# Combining live cell fluorescence imaging with *in situ* cryo electron tomography sheds light on the septation process in *Deinococcus radiodurans*

**DOI:** 10.1101/2024.11.18.624142

**Authors:** L. Gaifas, J.P. Kleman, F. Lacroix, E. Schexnaydre, J. Trouve, C. Morlot, L. Sandblad, I. Gutsche, J. Timmins

## Abstract

Cell division is a fundamental biological process that allows a single mother cell to produce two daughter cells. In bacteria, different modes of cell division have been reported that are notably associated with distinctive cell shapes, but in all cases, division involves a step of septation, corresponding to the growth of a new dividing cell wall, followed by splitting of the two daughter cells. The radiation-resistant *Deinococcus radiodurans* is a spherical bacterium protected by a thick and unusual cell envelope. It has been reported to divide using a distinctive mode of septation in which two septa originating from opposite sides of the cell progress with a flat leading edge until meeting and fusing at mid-cell. In the present study, we have combined conventional and super-resolution fluorescence microscopy of live bacteria with *in situ* cryogenic electron tomography of bacterial lamellae to investigate the septation process in *D. radiodurans*. This work provides important insight into (i) the complex architecture of the cell envelope of this bacterium, (ii) the ‘sliding doors’ septation process and (iii) the molecular mechanisms underlying septal growth and closure.

## INTRODUCTION

Bacterial cell division occurs through binary fission^1^ and involves two major steps: (i) septation, corresponding to the inward growth of a new dividing cell wall across the cell, known as septum or cross-wall, and (ii) separation or splitting of the two daughter cells notably through the action of cell wall hydrolases. These two steps can be either concomitant or dissociated in time. When splitting is sufficiently delayed compared to septation, the cross-wall advances centripetally like a closing iris until full closure. In this ‘septation’ mode of division, a septum can be observed at the division site. When septal synthesis and cleavage proceed at a similar rate, the division site undergoes an apparent ‘constriction’, with a reduction of its diameter without visible septum formation. Depending on the dynamics of septal wall synthesis and cleavage, bacteria have been proposed to divide either by septation, constriction, or a combination of these two processes^2–4^.

Bacterial cell division is tightly coupled to the synthesis and remodeling of the cell envelope, whose composition defines the classification between Gram-positive (or monoderm) and Gram-negative (or diderm) bacteria. In all bacteria, the cell envelope includes a common and essential constituent called the peptidoglycan (PG) that forms a cross-linked mesh-like structure, which confers cell shape and protects the cell from turgor pressure^5^. Gram-positive bacteria that are missing a second outer membrane bilayer usually possess a thick and multilayered PG, while Gram-negative bacteria have a thin predominantly single-layered PG^5^. Until recently, most Gram-negative bacteria were reported to divide through ‘constriction’. However, cryo-electron tomography (cryo-ET) studies have revealed that gonococci and *Escherichia coli* actually use a combination of ‘constriction’ and ‘septation’ to divide^6,7^. Similarly, in the Gram-positive bacterium, *Streptococcus pneumoniae*, both modes of cell division have been reported to be at play^8^. In contrast, many Gram-positive bacteria, including *Staphylococcus aureus* or *Bacillus subtilis*, divide through complete septation, with a septum composed of two adjacent cross-walls separated by a low density region^9–13^. In this mode of division, the diameter of the mother cell at the site of division is unaffected until splitting of the daughter cells. This splitting step can be slow and gradual as in *B. subtilis*^14^, or instead very fast through a ‘popping’ mechanism as in *S. aureus* and actinobacteria^15–17^.

These distinct modes of cell division rely on both common and species-specific molecular mechanisms and division factors. In most bacteria, cell division begins with the assembly at the division site of the highly conserved FtsZ protein into an annular structure known as the Z-ring on the cytoplasmic side of the inner membrane. This Z-ring then acts as a scaffold for the recruitment of several membrane-associated division and cell wall synthesis factors (including the penicillin-binding proteins or PBPs) that together form the divisome^18–20^. After the divisome has assembled, the Z-ring constricts and PG is synthesized at the leading edge of the invaginating membrane, dividing the mother cell into two daughter cells. In bacteria, two distinct PG synthesis machineries, the divisome and the elongasome, are responsible for septal and peripheral (or lateral) cell wall synthesis respectively^21^. In rod-shaped bacteria, the elongasome is organized by the actin-like MreB protein, which directs the so-called ‘lateral’ PG synthesis in the cylindrical region of the cell^22^. In cocci and ovococci, which are missing MreB, both the elongasome and divisome are recruited at mid-cell by FtsZ^3^. Recent studies making use of single-molecule localization microscopy (SMLM) and 3D structural illumination microscopy (3D-SIM), two super-resolution techniques that provide enhanced spatial resolution, have, however, revealed that despite being located at the same site in these bacteria, the divisome and elongasome nonetheless specifically drive septation and cell elongation processes, respectively^3,8,23–25^. While the divisome machinery synthesizes the septum, the elongasome synthesizes peripheral PG on either side of the division site, contributing to slight cell elongation.

The cell envelope composition of the spherical bacterium, *Deinococcus radiodurans*, known for its outstanding resistance to DNA-damaging agents^26–28^, remains a matter of debate. It is known to possess an unusual cell wall that stains Gram-positive, but yet is composed of both an inner and an outer plasma membrane interspersed by a thick multilayered region. The septation process in *D. radiodurans* is also unlike that of other cocci. Based on electron micrographs of freeze-cleaved *D. radiodurans* cells, Murray *et. al.* reported that division is not initiated from the whole circumference of the cell, but instead from two discrete regions on opposite sides of the cell. Two septa with a flat leading edge then progress across the cell until they fuse at mid-cell to close the slit^13^. This observation was more recently supported by 3D confocal microscopy imaging of Nile Red labeled *D. radiodurans* cell membranes^29^.

Exposure of *D. radiodurans* to high doses of radiation causes significant damage to the genome and immediate cell cycle arrest^30^, suggesting that cell growth and division in this organism are tightly regulated and coordinated with genome maintenance. *D. radioduran*s cells divide in two alternative perpendicular planes^13,31^ and their cell cycle can be classified into 6 phases (Figure 1A). Starting from an elliptical and largely symmetric pair of cells (or diad) in Phase 1, the cells grow and septal closure progresses until tetrads are formed in Phase 6, which, in exponential phase, are very short-lived and rapidly split into two diads to initiate a new cell cycle^29^. As a result, in this organism, the two major steps of cell division, *i.e.,* (i) septation and (ii) splitting of the daughter cells, actually occur in separate cell cycles, with septal closure taking place in cycle *n* and splitting of the cells in cycle *n+1* (Figure 1A). This temporal separation makes *D. radiodurans* particularly well suited as a model to study septation.

**Figure 1:**
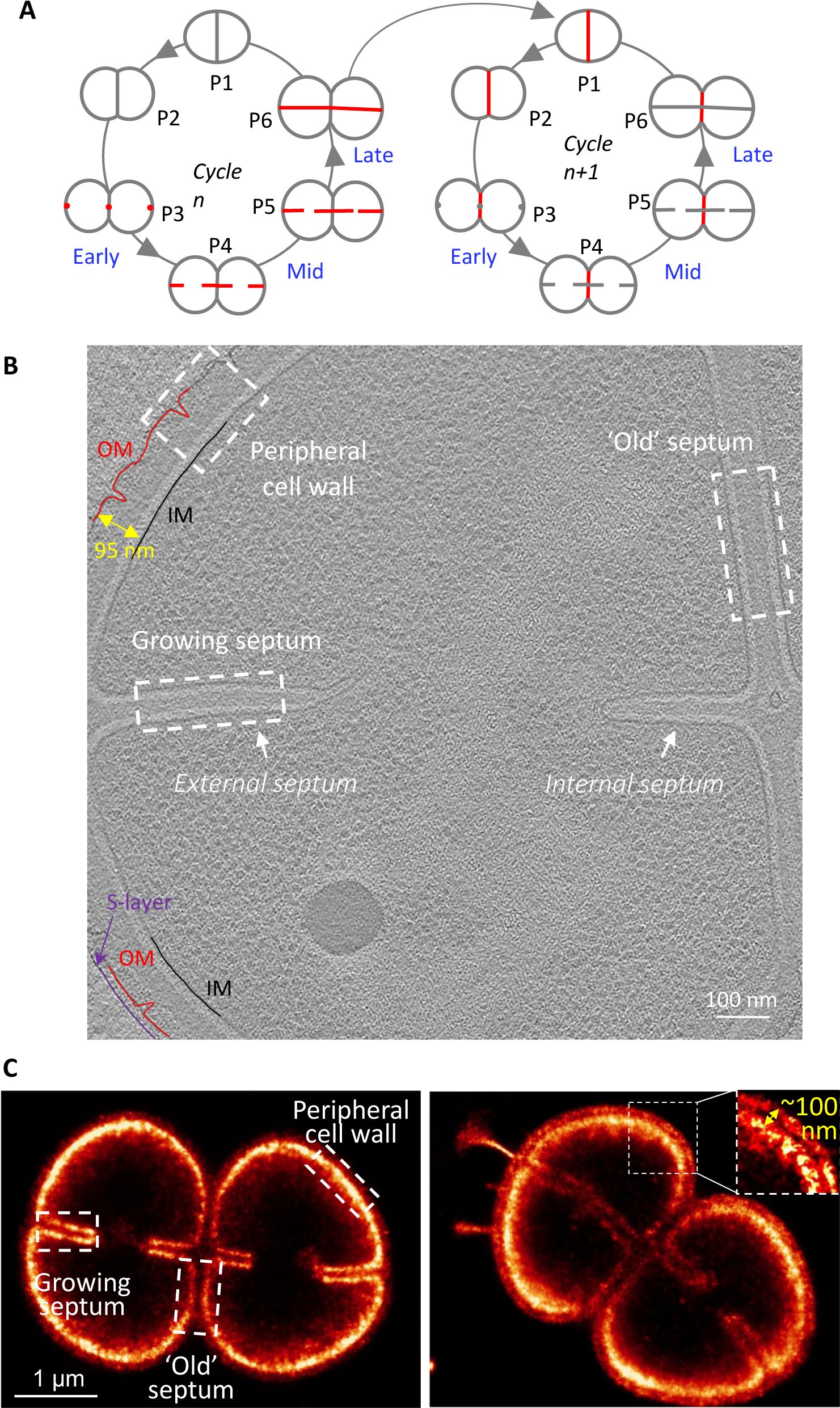
Structures of exponentially growing *D. radiodurans* cell walls during cell division. (A) Schematic diagram of exponentially growing and dividing *D. radiodurans*. Transition from a diad (two-cell unit) to a tetrad (four-cell unit) involves 6 distinct phases (P1-P6). The septation process occurs during phases P3 to P6 of cycle *n* (left, red), while the splitting of this newly synthesized cell wall only occurs in the following cell cycle, cycle *n+1* (right, red). Early-, mid- and late-phases of septation are annotated in blue. (B) Central slice of a typical cryo-electron tomogram of an actively dividing wild-type *D. radiodurans* diad. Three different types of cell envelope regions can be seen in this image: (i) the multilayered peripheral region composed notably of the inner (IM, black line) and outer (OM, red line) membranes and a discontinuous S-layer (purple line), (ii) the ‘old’ septum dividing the two cells composing the diad unit, and (iii) the ‘new’ growing septa originating from opposite sides of the cell. The mean distance between the two membranes in the peripheral region was determined to be 95 nm. The light grey density in the center of the cell corresponds to the nucleoid, the darker spherical form is a stress granule and the small dark densities distributed throughout the cytoplasm (and excluded from the nucleoid) are ribosomes. (C) Two examples of super-resolved PAINT images of Nile Red stained wild-type *D. radiodurans* diads. As in (A), three distinct regions of the cell envelope can be distinguished. On a few occasions, the two lipid bilayers of the peripheral cell envelope could be resolved (as in the right image). The mean distance between these two layers was determined to be 100 nm in good agreement with the 95 nm measured on the tomograms.

In the present study, we have combined conventional and SMLM microscopy of live cells with *in situ* cryo-ET of bacterial lamellae obtained by cryo-focused ion beam (FIB) milling, to image dividing *D. radiodurans* cells. This work unveils the different layers of the cell envelope at various stages of the division process. We unambiguously demonstrate that septation of *D. radiodurans* proceeds by a ‘sliding doors’ mechanism, in which the cross-wall grows through PG synthesis at the initially flat and progressively curved leading edge of the two opposing septa until membrane fusion occurs, first at the cell periphery and then all the way across the diameter of the cell in a zip-like mechanism. Using a fluorescent D-Ala probe that is integrated into newly synthesized PG, we further show that PG assembly in *D. radiodurans* occurs in both septal regions and throughout the whole periphery of the diad, and involves two distinct machineries for septal and peripheral cell wall synthesis, with the former being fully inhibited by ampicillin treatment. To our surprise, membranous protrusions were frequently observed in our tomograms at the tips of the septa at early stages of the septation process. We propose that these remarkable structures may constitute a preformed dual membrane layer and that PG synthesis in between these two lipid bilayers progressively fills, thickens and rigidifies the structure of the growing septa. Finally, this rigidification step appears to be associated with the assembly of FtsZ filaments at the tips of the growing septa.

## RESULTS

### Composition, structure and maturation of D. radiodurans cell envelope

To decipher the structure and composition of the unusual cell envelope of *D. radiodurans*, we have combined SMLM on live cells with cryo-ET on cryo-FIB-milled cell lamellae (Figure 1B-C). Three distinct cell envelope regions can be observed in our cryo-ET data (Figure 1B). The peripheral, fully matured, cell envelope is the thickest. The new growing septum is on the contrary the thinnest, and the ‘old’ septum (synthesized during the previous cell cycle) located between the two cells composing a typical *D. radiodurans* diad, corresponds to an intermediate stage of cell wall maturation (Figure 1B-C).

The peripheral cell envelope bears two lipid bilayers that can be seen as dark lines in the cryo-ET data (Figure 1B) and could be distinguished on a few occasions in the super-resolved PAINT images of Nile Red stained *D. radiodurans* (Figure 1C, right panel). The distance between the inner (IM) and outer (OM) cell membranes was in good agreement between the two techniques and was found to be ∼100 nm (Figure 1B-C). A more in-depth analysis of the cell envelope composition was facilitated by the calculation of straightened cell wall projections of the cryo-ET images using a recently developed tool, blik^32^. Density profiles of the peripheral envelope revealed that this region is composed of 5 distinct layers: (i) the IM, (ii) a low-density periplasmic space, (iii) a high-density PG layer, (iv) an intermediate layer previously described as the SlpA layer^33^ and (v) the OM (Figure 1B and 2A). In addition, in several tomograms, an additional S-layer was seen as the outermost layer of this complex cell envelope (Figure 1B, 2A and SI Fig. S1-S2). A distinctive white line was observed separating the PG layer from the SlpA layer, and V-shaped invaginations of the OM were seen regularly along the peripheral cell envelope. Measurements made on numerous tomograms (examples of which are provided in SI Fig. S1 and S2) allowed us to precisely define the architecture of this cell envelope region. When present, the S-layer was typically located at 17.9 ± 0.8 nm (n=5) above the OM. The SlpA layer was found to be 34.9 ± 1.5 nm (n=16) in thickness in good agreement with the estimated dimensions of the SlpA complex that stretches across this layer^33^. The PG layer was found to be 43.9 ± 4.8 nm (n=16) in thickness and located approximately 14.8 ± 3.2 nm (n=16) above the IM, with a low-density region nestled in between these two essential layers. Interestingly, these measurements revealed that the IM bilayer was significantly thicker than the OM (5.8 ± 0.5 vs 5.0 ± 0.5 nm; n=16), suggesting distinct lipid compositions (Figure 2B).

**Figure 2:**
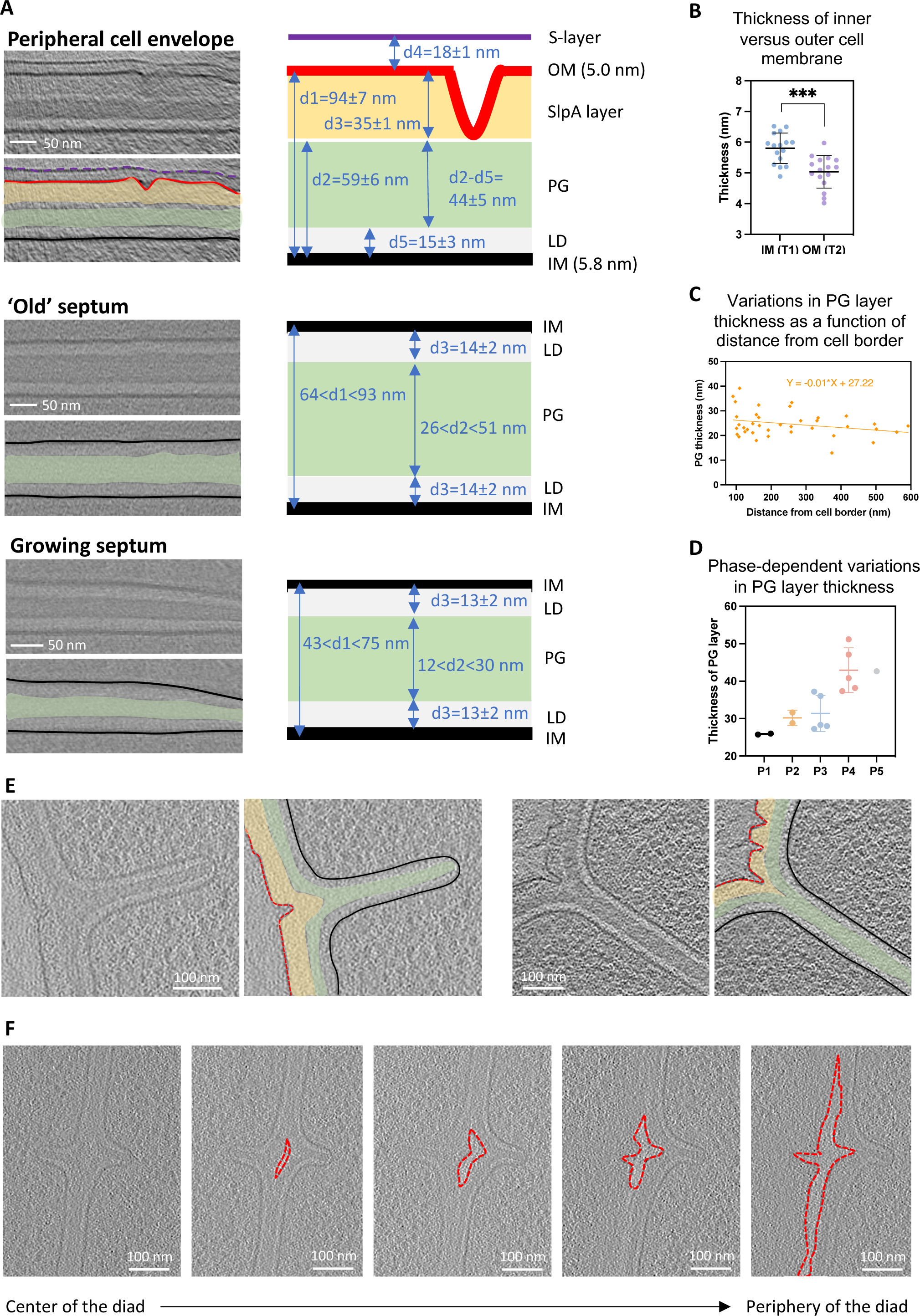
In-depth analysis of the architecture and composition of the cell envelope of *D. radiodurans*. (A) Straightened views of the peripheral region (Top), ‘old’ (middle) and growing (bottom) septa used to measure the thickness of the various layers, shown schematically to the right of the images (SI Fig. S10 for more details regarding the measurements). Left: Top panels show a straightened region and lower panels show the same region with segmentations of the various layers. The IM is highlighted in black, the OM in red, the S-layer in purple, the SlpA layer in yellow, the peptidoglycan (PG) in green and the low-density periplasmic space (LD) in light grey. (B) Mean and standard deviation of the thickness of the inner (IM) and outer (OM) membranes within the peripheral cell envelope. Individual points correspond to the mean thickness calculated from measures made on individual tomograms (N = 16). Horizontal and vertical bars correspond respectively to the mean and standard deviation of the mean thickness of each bilayer. Statistical test: Welch’s t-test. *** p=0.0002. (C) Plot illustrating the variation in PG thickness as a function of the distance from the root of the growing septa. Individual points correspond to single measurements made at different positions along the septa. (D) Plot illustrating the phase-dependent thickness of the PG layer in the ‘old’ septum (d2 in SI Fig. S10). Individual points correspond to the mean thickness calculated from measures made on individual tomograms (N < 6 for each phase). Horizontal and vertical bars correspond respectively to the mean and standard deviation of the mean PG thickness for each phase. (E) Two illustrations of junctions between the peripheral cell envelope and the growing septa. Left panels show 2D slices of a tomogram and right panels show the same regions with segmentations of the various layers (colors as in A). (F) 2D slices extracted from different Z values of a given tomogram, illustrating the progressive splitting of daughter cells that starts at the cell periphery of the diad (right) and then progresses inwards towards the center of the diad (left). The OM is delineated in red.

A similar procedure was used to analyze the composition of the growing and ‘old’ septa (Figure 1B and 2A). The outermost layers of the peripheral cell envelope (S-layer, OM and SlpA layer) were missing in these regions, and instead septa were composed of a single continuous PG layer sandwiched between two low-density periplasmic layers and bordered on either side by the IM (Figure 2A). These three layers were continuous with those of the peripheral cell envelope. While the IM bilayer and low-density periplasmic space showed similar thicknesses in these two locations, the PG layer displayed significant variations, ranging from 26 to 51 nm in the ‘old’ septa and as low as 12 nm in the growing septa. In both septal regions, the PG layer was typically thicker at the cell periphery than at the leading edge of the septum (Figure 2C) or the center of the diad. Moreover, in the ‘old’ septa, the mean thickness of the PG layer was also found to vary substantially as a function of the phase of the cell cycle (Figure 2D), suggesting a progressive thickening of this layer until the splitting of the daughter cells.

At the junction between the peripheral cell envelope and the root of the growing septum, we observed that the PG and low-density periplasmic layers followed the IM, while the outermost layers (SlpA layer and OM) were restricted to the peripheral cell envelope and were often seen to form V-shaped invaginations (Figure 2E and SI Fig. S3A for more examples). 3D visualization of the tomograms revealed that these invaginated outer layers extended inwards towards the center of the diad during the splitting of the two daughter cells, forming bubble-like structures (Figure 2E-F and SI Fig. S3B for more examples).

### Structure of septal tips

A close inspection of the tomograms revealed that the tips of the growing septa exhibited particular structures. A majority of septa (40 of the 64 septa visible in our tomograms) were slightly tapered at their tips and the low-density periplasmic space was significantly thinner in these regions, bringing the PG layer very close to (in some cases even contacting) the IM (Figure 3A). Strikingly, in nearly 40% of the growing septa, membrane protrusions were observed at their tips (Figure 3B-C and SI Fig. S4 for more examples). These structures, mostly observed in cells at early stages of septation, appear to be composed solely of the low-density layer delimited on either side by the IM, and adopt either outstretched tube-like structures or more circular loop-like arrangements. In all cases, they appear to be very flexible, probably as a result of the absence of PG to rigidify the protrusion. This intrinsic flexibility may explain why the tips of the growing septa were often not observed in the super-resolved images of Nile Red-labelled *D. radiodurans* cells (Figure 3D, left panel). Instead, in these images, open ends were observed at the leading edge of the growing septa. This suggests that either the Nile Red dye poorly labelled these highly curved membrane bilayers (likely exhibiting altered membrane fluidity that is known to affect Nile Red staining^34^) or that the tips were very mobile and not captured in live imaging experiments; both of these phenomena may also be at play. On a few occasions, we did nonetheless observe poorly defined Nile Red labelling at the tips of growing septa that may correspond to such flexible membrane protrusions (Figure 3D, right panel and Figure 1C, right panel).

**Figure 3:**
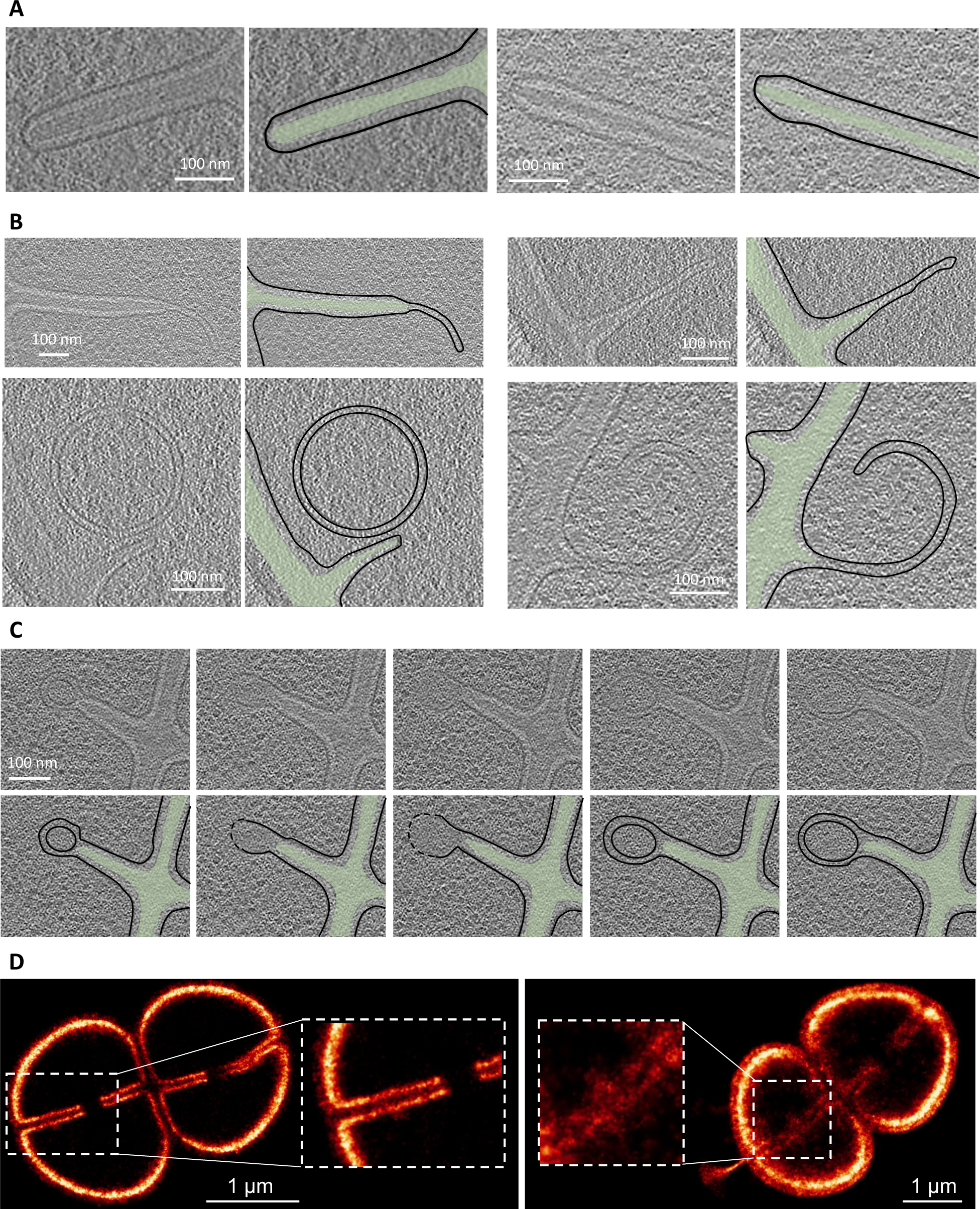
Close-up view of septal tips. (A-B) Examples of typical septa tip morphologies: (A) tapered leading edge with a well-defined and rigid PG layer (green) stretching almost to the IM (black) and (B) tubular (top) and curved (bottom) membrane protrusions extending the growing septa. Left panels show 2D slices of a tomogram and right panels show the same regions with segmentations of the various layers (colors as in Fig. 2A). (C) 2D slices extracted at different Z values from a given tomogram illustrating the change in size and shape of these membrane protrusions as a function of the position in the tomogram. Top panels show 2D slices and lower panels show the same regions with segmentations of the PG and IM layers (colors as in Fig. 2A). (D) Close-up views of the leading edges of growing septa captured by PAINT microscopy of Nile Red stained wild-type *D. radiodurans* diads in the process of dividing. Left image: septal tips appear to be open-ended with no fluorescence signal for the highly curved membrane located at the leading edge of these closing septa. Right image: blurry septal tips are visible pointing in opposite directions. These structures may correspond to the membrane protrusions observed in the tomograms (B, C).

### Septation through a ‘sliding door’ mechanism

Using timelapse 3D video confocal microscopy of Nile Red stained *D. radiodurans* bacteria immobilized in various orientations on agarose pads (SI Fig. S5), we followed the division process in live cells (SI video 1). Septation was found to proceed in several steps. First, two septa originating from opposite sides of the cell grow inwards with a flat leading edge (Figure 4). As the septa grow, their leading edge progressively becomes more curved. Finally, when the two septa come close to each other, fusion starts first at the periphery of the cell, forming a cat’s eye structure, and then rapidly proceeds through a zipping mechanism towards the cell center. 3D PAINT images of Nile Red labeled *D. radiodurans* cells confirmed these observations, allowing to capture snapshots of dividing cells exhibiting septa with flat or slightly curved leading edges embracing a central slit that stretches across the cell (Figure 4A). Kymographs of individual septation events were extracted from the live cell confocal acquisitions to probe the kinetics of septal closure (Figure 4B). These revealed that septation is a linear process in which the external and internal septa (Figure 1B) grow respectively at rates of 7.3 ± 0.5 nm.min^-1^ and 4.1 ± 0.7 nm.min^-1^ until full closure. Interestingly, the external septum not only grows at a faster rate than the internal septum, as can be seen from the asymmetric V-shaped structure of the kymographs, it also starts growing ahead of the internal septum (Δt in Figure 4B). This asymmetry in the growth of the internal and external septa was also clearly visible in the tomograms, capturing bacteria at various stages of the division process (Figure 4C-D). This is likely explained by the diad configuration of *D. radiodurans* cells and their expansion that is concomitant with the septation process (Figure 1A). Indeed, the external septum must grow further than the internal one to reach the site of fusion located on the plane that intersects the point of maximal curvature of *D. radiodurans* cells. The flat or slightly curved leading edges of the growing septa were also observed in the tomograms (Figure 4C).

**Figure 4:**
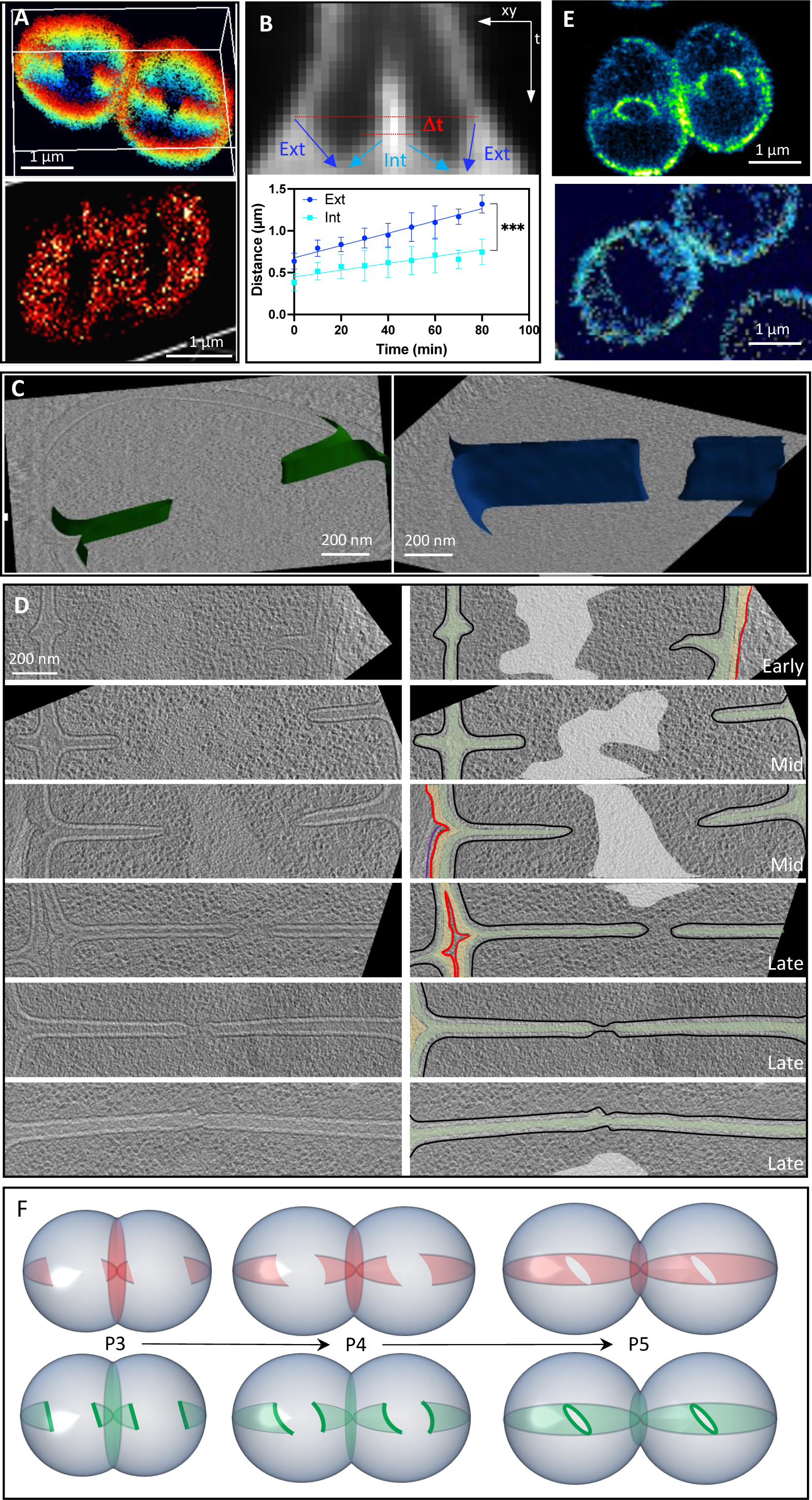
Septation through a ‘sliding doors’ mechanism. (A) 3D super-resolved PAINT image of a Nile Red stained wild-type *D. radiodurans* diad. Top panel: the 3D volume was obtained by combining 2 stacks of images collected at different focus heights. The flat leading edge is very clear in the left cell while the right cell, which is more advanced in its division process, illustrates the formation of a cat’s eye structure stretching across the cell. This structure is also visible in the lower panel, corresponding to a side view of a labeled cell. (B) Kymograph analysis of the septation process in Nile Red stained *D. radiodurans*. Top panel: example of a typical kymograph obtained for a dividing diad. The coordinates of the borders of the fluorescent signal were used to determine the length of the external (dark blue) and internal (light blue) septa as a function of time. The lag time (Δt) between the start of growth for the external septa versus the internal septa was also measured. Lower panel: Plot illustrating the linear growth of the external (dark blue) and internal (light blue) septa as a function of time. Data points correspond to the mean lengths of measured septa (4 < N < 12) at each time point. Error bars correspond to the standard deviation. The external septa were found to grow at a significantly higher rate than the internal septa. *** (p-value: 0.0004). (C) Examples of segmented septa in two tomograms, illustrating the flat edge of the growing septa (left panel, green) that progressively becomes more curved (right panel, blue) as the division process advances. (D) 2D slices extracted from various tomograms illustrating different stages of the septation process (Early, Mid-or Late) in *D. radiodurans*. Left panels show 2D slices of a tomogram and right panels show the same regions with segmentations of the various layers (colors as in Fig. 2A). The light grey area in the right panels corresponds to the nucleoid. (E) 3D super-resolved dSTORM image of PG-labeled (through incorporation of azido-D-Alanine) wild-type *D. radiodurans* diads. Top panel: PG synthesis occurs both in the septa and in the peripheral cell wall, with a strong PG synthesis activity detected at the leading edge of the growing septa (intense ring and arch). Lower panel: side view of a PG-labeled *D. radiodurans* diad, illustrating the incorporation throughout the septal region and the formation of the cat’s eye structure similar to that observed in Nile Red stained cells (A). (F) Schematic model of the ‘sliding door’ mechanism of septation in *D. radiodurans*. The top panels illustrate the growth of the membrane (red), while the lower panels show the closely-related growth of the PG layer (green) and the strong PG synthesis activity at the leading edge of the septa.

In addition to staining the cell membrane with Nile Red, we also labeled the PG by growing *D. radiodurans* cells in the presence of azido-D-alanine (aDA) during a short (10 min) ‘pulse labeling’ period. The incorporated aDA moiety was subsequently conjugated to fluorescent probes suitable for either confocal or dSTORM imaging^8,35^ by copper-free click chemistry (SI Fig. S6). The aDA pulse-labeling pattern was very similar to that of Nile Red, with efficient incorporation in both the growing and ‘old’ septal regions, as well as in the peripheral cell envelope (Figure 4E and SI Fig. S7), as observed previously using fluorescently labeled D-alanine^29^. In both the 3D dSTORM and confocal images, the observation of tilted cells offering a front or side view (as opposed to the more common top view) of the septation plane further revealed that PG labeling is enriched at the leading edge of the growing septa, suggesting it corresponds to the major site of septal PG synthesis in these cells (Figure 4E-F and SI Fig. S7). This was particularly visible at late stages of the cell cycle (Figure 1A), either just before (Phase 5) or just after septum closure (Phase 6), where an intense band of labelled PG could be seen at the site of fusion (SI Fig. S7C). This finding was confirmed by performing a pulse-chase experiment in which aDA pulse-labeled cells were further grown for 45 min in the absence of the probe (SI Fig. S8A). In the absence of the chase, Nile Red and PG-labeling were largely superimposed. In contrast, after the chase, most Nile Red-stained septa showed incomplete PG labeling, indicating that septa having incorporated aDA at their leading edge during the pulse period were extended by unlabeled PG during the chase period. Altogether, these observations indicate that new PG is added to the existing layer in an inwards direction until the opposing septa are close enough to fuse (Figure 4F and SI Fig. S8B). On the contrary, peripheral PG labeling was largely unaffected by the chase, suggesting that aDA incorporation (and thus PG synthesis or remodeling) occurs throughout the periphery of the diads, leading only to a progressive dilution of the labeling intensity (SI Fig. S8B).

Close examination of the tomograms also revealed that the relative position and shape of the growing septa changed as a function of their length (Figure 4D). At early stages of the septation process, short septa were not always precisely facing each other and their tips were often bent and bearing membrane protrusions (Figure 3B-C and Figure 4D), while at later stages, septa were remarkably straight and fully aligned to ensure their fusion (Figure 4D). The presence of a thick PG layer in the growing septa appears to be a pre-requisite for the formation of these straight and well-aligned septa, indicating that PG synthesis may provide the necessary rigidity for septal closure. This final fusion step was captured in two of the tomograms and was found to proceed first through fusion of the IM and subsequently through synthesis of PG to fill the gap and ‘*glue’* the two septa together (Figure 4D and SI Fig. S2). Taken together, these data allow us to propose a model of septation through a ‘sliding doors’ mechanism, which is illustrated schematically in Figure 4F.

### PG synthesis in the peripheral and septal regions are performed by distinct machineries

To better understand the molecular mechanisms underlying this unusual mode of septation, we treated *D. radiodurans* cultures with a β-lactam antibiotic, ampicillin, and observed the growth of untreated and treated Nile Red labeled cells by 3D confocal timelapse microscopy for a three-hour period (Figure 5A and SI videos 2 and 3). Ampicillin is known to bind to the active sites of certain PBPs, thereby inhibiting their enzymatic PG synthesis function^36^. To our surprise, septation, but not cell growth was arrested by ampicillin treatment (Figure 5A-C). Indeed, the growth of the cell perimeter was unaffected by this treatment during this three-hour period, while growth of both the internal and external septa were rapidly arrested (Figure 5B-C). After 1 hour, the septa even started to shorten, suggesting they may be undergoing disassembly and retraction (Figure 5C). As a result of this change in the balance between cell growth and septation, cells progressively became distorted (more elongated with a slight bulge at sites of division), as evidenced by the substantially increased diameter (d) to perimeter (p) ratio after 3 h of treatment with ampicillin (Figure 5D-E). The specific inhibition of septal growth by ampicillin suggests that the PBPs responsible for septal PG synthesis and maturation are distinct from those involved in peripheral PG synthesis (Figure 5F).

**Figure 5:**
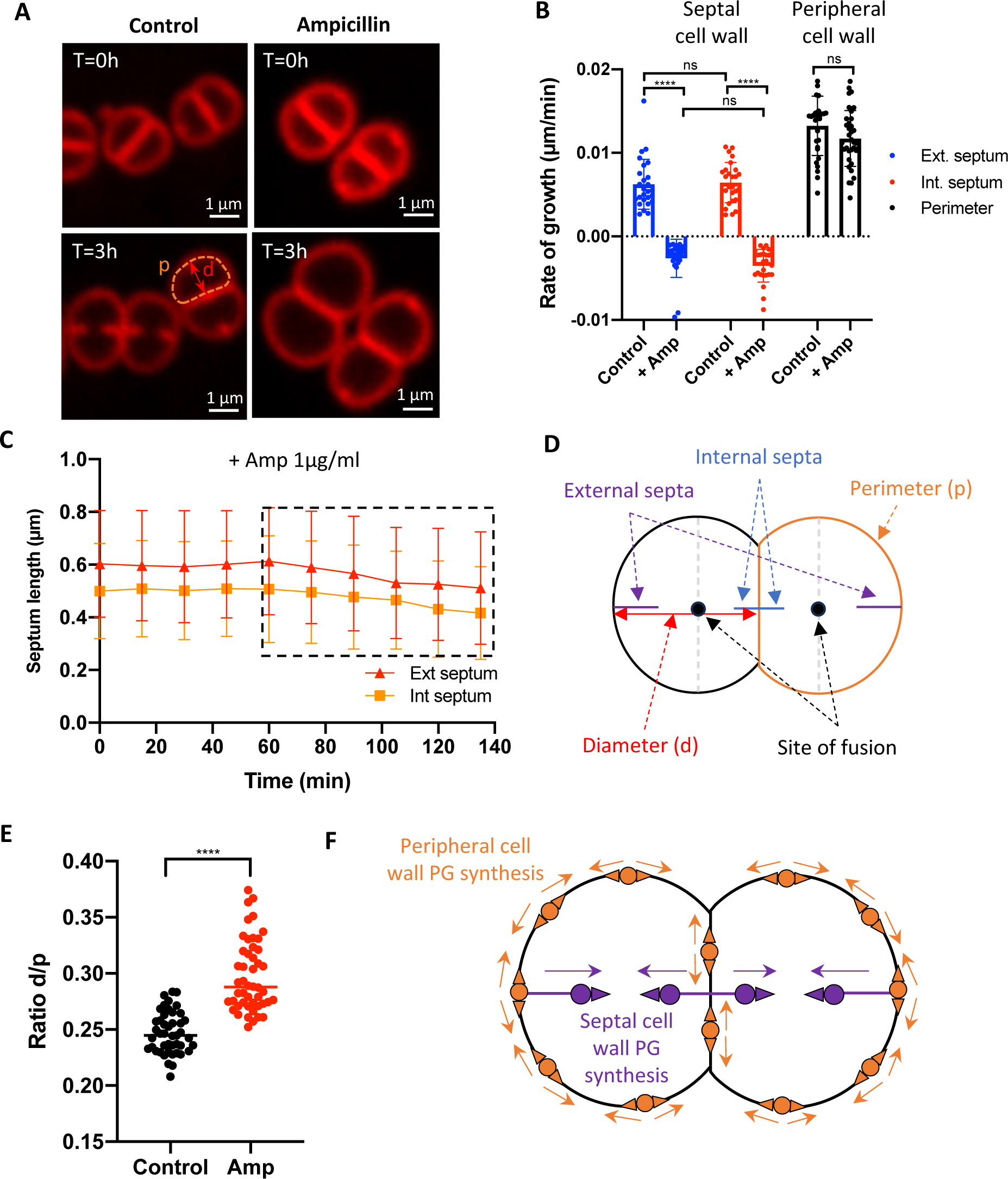
Distinct machineries are involved in septal and peripheral PG synthesis. (A) Examples of untreated (left) and ampicillin-treated (right) Nile Red stained *D. radiodurans* at the start (T = 0) and end (T = 3 h) of the time-lapse experiment (see SI videos 2 and 3). (B) Effects of ampicillin treatment on the growth rate of the external (blue) and internal (red) septa, and of the peripheral cell wall (black) derived from measurements of the cell perimeter (N > 25). Statistical test: One-way ANOVA. ns: non-significant, **** p-value < 0.0001. (C) Plot illustrating the inhibition of both the external (red) and internal (orange) septal growth by treatment with 1µg/mL ampicillin. After 1 h of treatment (designated with a dashed box), the lengths of the septa. (D) Schematic diagram of a *D. radiodurans* diad, illustrating the measurements performed on different features to probe the effect of ampicillin treatment. (E) Comparison of the diameter:perimeter (d/p) ratio of untreated (black) and ampicillin treated (t = 3 h) cells (red). Data points correspond to individual measurements made on N > 45 cells. Horizontal bars correspond to the mean values. Distortion of the cell morphology as a result of ampicillin treatment is particularly visible and is reflected in the marked increase in the d/p ratio in the presence of ampicillin (red). Statistical test: unpaired t-test. **** p-value < 0.0001. (F) Schematic model of the two PG synthesis machineries at play in *D. radiodurans*. Peripheral PG synthesis and remodeling (orange) that is insensitive to ampicillin is likely to occur throughout the peripheral envelope of the diads. In contrast, septal PG synthesis (purple) is very sensitive to ampicillin and is located mainly at the leading edge of the growing septa.

Next, we repeated the aDA pulse-labeling experiment on cells pre-grown for either 1 or 2h in the presence of ampicillin before aDA incorporation (SI Fig. S9). In agreement with our timelapse experiments, aDA was still incorporated in the peripheral region, albeit to a reduced level compared to untreated cells. Septal labeling, on the other hand, was restricted to bright dots (in 2D) or distinctive rings (in 3D) positioned at the division site (SI Fig. S9). In untreated cells, such PG-labeled rings were typically seen in ∼ 10% of cells, corresponding to cells at the onset of division. In ampicillin-treated cells, and more particularly in samples pre-grown for 2 h in the presence of the β-lactam antibiotic, over 30% of cells exhibited such PG-labeled rings (N > 300; SI Fig. S9B). Initial septal PG synthesis thus appears to be unaffected by ampicillin, while subsequent extension of the septa is arrested by this treatment. Distinct sets of PBPs may therefore be involved in the initiation and extension stages of septation (Figure 5F).

### FtsZ is present at the tips of septa

We investigated the location of FtsZ, one of the key players in bacterial cell division, in dividing *D. radiodurans* cells using both fluorescence microscopy and cryo-ET. Two strategies were used to fluorescently label FtsZ: (i) immunolabeling of an endogenously HA-tagged FtsZ or (ii) endogenous tagging of FtsZ with a photoconvertible fluorescent protein, mEos4B. The latter was compatible with live cell imaging, but this genetically modified strain of *D. radiodurans* showed altered cell morphology (forming many large cells with likely impaired septation) and only a small fraction of cells exhibited fluorescence signal and visible Z-rings (Figure 6A). In contrast, the strain expressing HA-tagged FtsZ showed no obvious morphological defects, and FtsZ could be detected by either confocal microscopy or dSTORM, following anti-HA antibody immunolabeling of fixed and permeabilized cells (Figure 6A-B). Both strategies indicated that *D. radiodurans* FtsZ forms ring- or oval-shaped structures, some of which appear to be incomplete and/or display weaker curvature in regions likely corresponding to septal leading edges. In support of these observations, dual labeling of FtsZ and the cell membrane showed that FtsZ locates to the leading edge of growing septa at all stages of septation, including very early stages when septa are not yet visible by fluorescence microscopy (Figure 6B).

**Figure 6:**
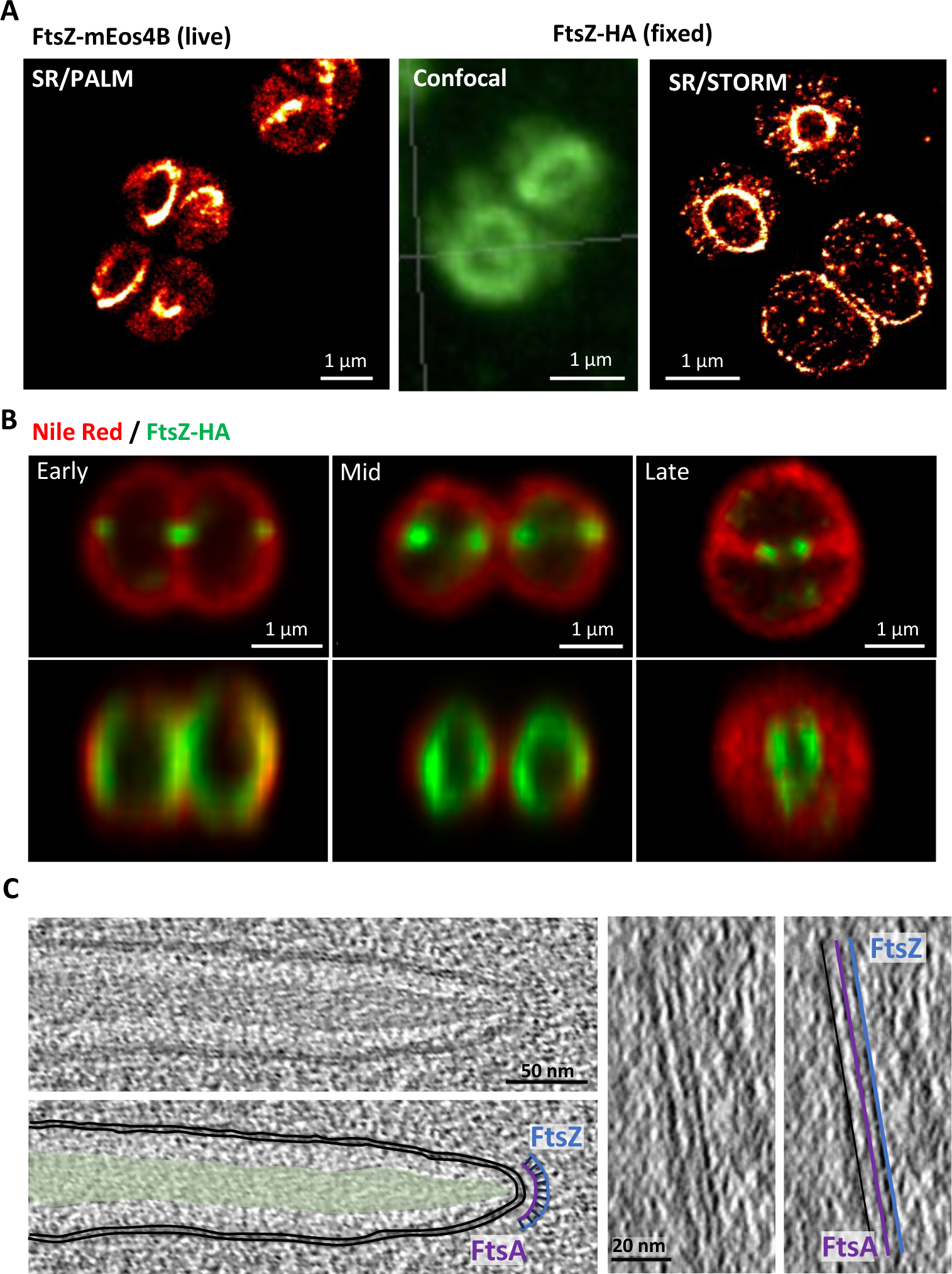
Localization of the key cell division factor, FtsZ, in *D. radiodurans*. (A) 3D fluorescence imaging of FtsZ in *D. radiodurans*. Left: 3D super-resolved PALM imaging of *D. radiodurans* cells expressing FtsZ-mEos4B, which forms well-defined ring-like structures. Middle and right: 3D confocal images (middle) and super-resolved dSTORM images (right) of immunolabeled *D. radiodurans* cells expressing FtsZ-HA. (B) Two-color labeling of *D. radiodurans* cells expressing FtsZ-HA (green) and stained with Nile Red (red). From left to right: illustrations of different stages of the division process (early, mid- and late). At all stages, FtsZ localizes to the leading edge of the septa. Top panels: top views, lower panels: side views. (C) Left: 2D slice of a typical tomogram of *D. radiodurans* illustrating the double-arched structure of FtsA (purple) and FtsZ (blue) located ∼ 15 nm away from the IM (black) on the cytoplasmic side of the growing septum. Inter-filament distance was estimated to be ∼ 5 nm. The top panel shows a 2D slice through a tomogram and the lower panel the same region with segmentations of the various layers (colors as in Fig. 2A) and the double-arched structure. Right: side view of the FtsA and FtsZ filaments highlighted in the right panel in purple and blue respectively.

In several tomograms, a double-arched structure was seen in the cytoplasm at ∼ 15 nm from the IM border of the septal leading edge, which likely corresponds to FtsZ (outer arch) and its cellular partner, the membrane-bound FtsA^12^ (inner arch; Figure 6C). These two arches follow the curvature of the septal tip, with the outer FtsZ arch typically between 20 and 50 nm in length. As in our fluorescence microscopy data, these structures were observed at all stages of the septation process from budding septa all the way to almost fusing septa. In the side projections of the tomograms, FtsZ was observed to form long, straight filaments along the flat leading edge of the closing septa, with an inter-filament distance of approximately 5 nm, in good agreement with previous reports of *in situ* FtsZ filaments^37,38^ (Figure 6C). Interestingly, when membrane protrusions were present at septal tips, FtsZ was not observed on these protrusions, but was occasionally seen in Z-slices either above or below these structures, which may explain some of the discontinuous ring structures observed by fluorescence microscopy (Figure 6A-B).

## DISCUSSION

In this study, we have combined live conventional and super-resolution fluorescence microscopy with *in situ* cryo-ET imaging of *D. radiodurans* to follow the process of septation in this unusual, spherical bacterium. This work provides important insight into (i) the complex architecture of its cell envelope along the cell cycle, (ii) its distinct mode of septation, involving a ‘sliding doors’ mechanism based on the growth of two septa originating from opposite sides of the cell, and (iii) the sequence of events underlying membrane and PG synthesis during septal growth. Furthermore, it starts to unveil the key role of FtsZ in this division process.

*D. radiodurans* possesses a unique cell envelope including an outer S-layer, which has been the object of numerous studies over the past decades^33,39–47^. Like other members of the *Deinococcus-Thermus* phylum, *D. radiodurans* exhibits features of both Gram-positive and Gram-negative bacteria, and may be considered as a primitive form of diderm bacterium^4^. *D. radiodurans* is indeed lacking lipopolysaccharides commonly found in Gram-negative bacteria, possesses a thick PG layer (35-55 nm) typical of Gram-positive bacteria and yet its cell envelope is composed of two membrane bilayers characteristic of Gram-negative bacteria^48^. The exact composition and structure of this unusual cell envelope has been the object of much controversy in recent years, notably regarding the outer layers linking the PG to the outermost S-layer hexagonal lattice structure. Our in-depth cryo-ET analysis of the complex architecture of *D. radiodurans* cell envelope fully supports the model recently proposed by Bharat and colleagues, in which the whole peripheral cell envelope is ∼ 100 nm in thickness and composed of two membranes in between which can be found a thin periplasmic space and the thicker PG and SlpA layers^33^. The S-layer forms an additional coat located ∼ 18 nm above the OM^41^. The SlpA layer takes its name from the major protein constituent of this layer, the SlpA protein, a trimeric porin-like protein that is embedded in the OM and stretches across the SlpA layer to connect to the PG layer via a long coiled-coil region^33^. The predicted length of this assembly (∼ 28-29 nm) is in good agreement with our measured thickness of the SlpA layer (35 nm). Interestingly, in our tomograms, we observe a distinctive white line between the PG and SlpA layers that may correspond to the sites at which the flexible N-terminal SLH domain of SlpA attaches to the PG layer.

In *D. radiodurans*, daughter cell separation driven by the activity of autolysins^49^ (four of which have been identified in *D. radiodurans*^50^) is uncoupled from septation, with these two phenomena occurring in successive cell cycles. Furthermore, we show that three cell envelope regions can be distinguished in dividing *D. radiodurans*, displaying different compositions and characteristics that reflect their stage of maturation. Contrary to *S. aureus*, in which two distinct PG layers (one for each daughter cell) are readily observed in growing septa^6,10^, all *D.* radiodurans septa are composed of a single PG layer, which thickens during septal growth and even after closure of the septal slit. The SlpA and OM layers of the outer envelope are only added at a late division stage, when the splitting of the cells is initiated. Different from *E. coli*, in which the full cell envelope is synthesized within one division cycle^6^, the assembly of the cell envelope in *D. radiodurans* therefore encompasses two complete cell cycles. The temporal dissociation between the synthesis of the inner layers (IM and PG) and that of the outer layers (SlpA and OM), which is coordinated with septum splitting, may be required to allow incorporation of the abundant SlpA protein onto the PG layer and its attachment to the OM. These two additional layers efficiently protect *D. radiodurans* from its external environment and must therefore be completed before the splitting of the daughter cells. Our 3D cryo-ET data suggest that the splitting process is initiated from the periphery of the cell at sites of OM invagination and then moves inwards towards the center of the diad until full separation of the two daughter cells.

PG synthesis and its subsequent remodeling are key processes in bacterial septation. In *D. radiodurans*, PG synthesis occurs in both the septal and peripheral cell envelope regions^29^. Here, by observing aDA-labeled cells with high-resolution fluorescence microscopy techniques, we show that the leading edge of the septa constitutes the main site of septal PG synthesis in dividing *D. radiodurans* and that it proceeds in an inwards direction until the opposing septa meet and fuse. These growing septa are very often tapered with a thinner PG layer at the leading edge than at the root of the septum. This has also been reported in *S. aureus*^10,25,51^, suggesting that PG synthesis may not be restricted to the leading edge of the septa. As has been reported for the ovococci, *Lactococcus lactis*^24^ and *S. pneumoniae*^52,53^, we found that ampicillin treatment specifically impedes septation and not peripheral cell wall expansion needed for cell growth. In ovococci, this effect has been attributed to the specific inhibition of the class B PBP, PBP2x, that is part of the divisome and is involved in septation^24,52,54^. The genome of *D. radiodurans* encodes for two class A PBPs and one class B PBP, which is annotated as PBP2 (DR_1868). Although little is known so far about the respective roles of these PBPs in *D. radiodurans*, our observations suggest that *D. radiodurans* PBP2 may be the target of ampicillin and thus largely responsible for septal PG synthesis at the leading edge of the growing septa.

Whereas in most cocci and ovococci studied so far septation advances centripetally like a closing iris, with the present study, we provide evidence that septation in *D. radiodurans* is quite distinct. It proceeds via a ‘sliding doors’ mechanism, also described in an earlier study as a ‘septal curtain’^13^, in which the two septa originating from opposite sides of the cell grow inwards with a flat leading edge. As septation progresses, the leading edge becomes more curved and eventually, when the two septa come close to each other, fusion occurs and the septal disk is filled. Why *D. radiodurans* uses such a remarkable mode of septation remains to date a mystery. Physical constraints associated with the relatively large size (2-3 µm in diameter) of *D. radiodurans*, its cell morphology (diads and tetrads) or its mode of division in two alternating perpendicular planes may have contributed to the development of this unusual division mechanism.

With such a septation process, ensuring the opposing septa are correctly aligned and meet for septal fusion constitute major challenges for the bacteria. The latter is, in part at least, achieved by differential growth rates of the external and internal septa. Indeed, we observed that the septum originating from the external side of the diad grows at a faster rate and initiates its growth shortly before the internal septum originating from the ‘old’ septum to compensate for the longer distance needed for it to reach the site of fusion. This study also identifies two important factors that may constitute a physical link between the opposing septa: (i) the flexible membrane protrusions at the septal tips, and (ii) the FtsZ rings that encircle the division plane. These two mutually exclusive features are present at the leading edge of *D. radiodurans* septa. Membrane protrusions were detected in many instances at early stages of the septation process, adopting a variety of conformations. Interestingly, similar structures were reported in early electron microscopy studies of *D. radiodurans* ^31,55^ and other bacteria^56,57^, but were later considered as artefacts of chemical fixation procedures used for sample preparation^58–60^. In our case, these structures were observed *in situ* in near-native conditions (vitrified cells), strongly suggesting that they are not artefacts. Instead, we propose that these thin, mobile membrane extensions, missing a PG layer, may provide the necessary flexibility to facilitate the alignment of opposing septa during the initial steps of septation. In contrast, we found that the presence of FtsZ filaments at the septal tips was associated with more rigid and well aligned septa, suggesting that the assembly of the FtsZ ring may constitute a critical step in the proper alignment of opposing septa, possibly through a direct bridging of the two ‘sliding doors’. Importantly, FtsZ was only observed at the leading edge of septa containing a well-defined central PG layer that extended almost to the lipid bilayer. This suggests that the Z-ring may promote septal extension by recruiting the PG synthesis machinery to the septal tips, as has been reported for many other bacteria (recently reviewed in^62–64^). However, FtsZ may not be needed for septal initiation since aDA-labeled rings corresponding to early stages of septum synthesis were still observed in the presence of ampicillin, which is known to inhibit FtsZ-associated PBPs^61^.

Although FtsZ was never seen on the membrane protrusions themselves, in several tomograms FtsZ was observed on the leading edges of such septa in Z-slices located either above or below the protrusions. The leading edge of a given ‘sliding door’ is thus likely to be both dynamic and heterogeneous, composed of flexible regions bearing membrane protrusions and more rigid areas in which FtsZ can assemble to drive PG synthesis. We thus propose a model for *D. radiodurans* septation (Figure 7) in which septal PG synthesis is initiated by ampicillin-insensitive PG synthases, and then further extended by ampicillin-sensitive PBPs that are likely part of the divisome in a FtsZ-dependent manner. In this model, FtsZ-independent membrane synthesis would precede FtsZ-driven PG assembly to create membrane protrusions that would facilitate alignment of the opposing septal sites at early stages of the division process. The subsequent assembly of FtsZ filaments at the leading edge of the septa, followed by the formation of the Z-ring, would then act as a cue for the recruitment and activation of PG synthesis in order to progressively fill and rigidify the growing septa. Further studies will certainly be needed to decipher the precise details of this process and the specific role of FtsZ and other key players in this complex and intriguing septation of *D. radiodurans*.

**Figure 7:**
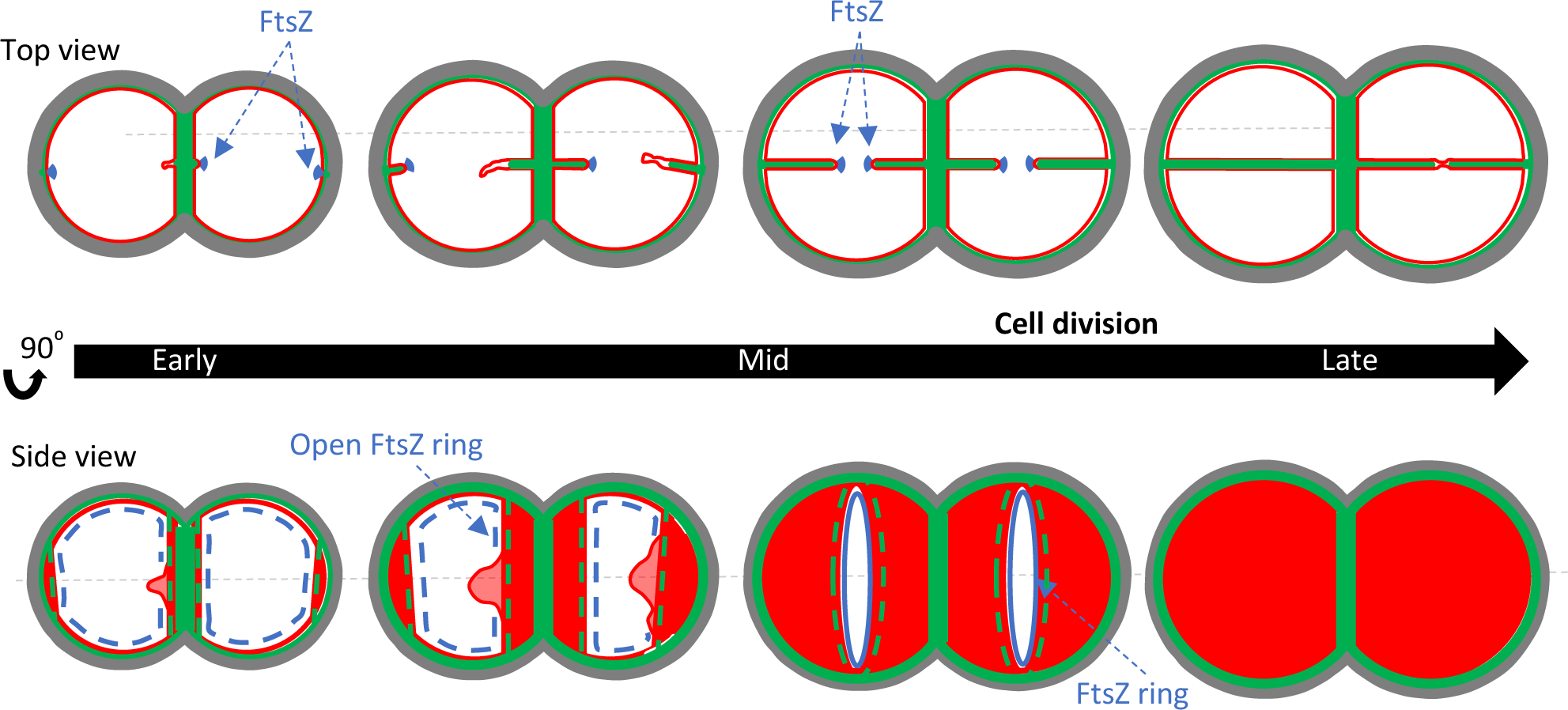
Schematic model of the septation process in *D. radiodurans*. Top and side views are illustrated at various stages of the division process. For simplicity, only the IM (red) and the PG layer (green) are highlighted. FtsZ is shown in blue with incomplete Z-rings as dashed lines and complete Z-rings as a full line. Flexible membrane protrusions are visible at early stages of the septation process, and are progressively filled with PG, providing the necessary rigidity to achieve a solid and well-aligned cross-wall. FtsZ plays a key role in targeting the ampicillin-sensitive PG synthesis machinery to the leading edge of the growing septa and may also facilitate the proper alignment of opposing septa by bridging them via Z-ring formation.

## MATERIALS & METHODS

### Bacterial cultures

*D. radiodurans* strains used in this study are listed in Table S1. All strains were derivatives of the wild-type strain R1 ATCC 13939 (DR^WT^). The genetically engineered strain of *D. radiodurans* expressing FtsZ fused to mEos4B (DR-*FtsZ-mEos4B*) was obtained by the tripartite ligation method as described recently^65^. A synthetic gene encoding mEos4B was amplified together with the kanamycin resistance cassette by PCR as were the regions (∼ 500 bp) flanking the insertion site (3’ end of *ftsZ* gene and region immediately downstream of the *ftsZ* gene) using oligonucleotides listed in Table S2. After restriction digestion the three fragments were ligated together and transformed into *D. radiodurans*. Transformants were selected on Tryptone-Glucose-Yeast extract (TGY) agar plates containing 6 µg/mL kanamycin, leading to allelic replacement on one genome copy. Because *D. radiodurans* is multigenomic, the transformant colonies were streaked three times successively on selective medium to ensure that all copies of the genome had incorporated the foreign DNA. This was then confirmed by PCR analysis and DNA sequencing. *D. radiodurans* cells were grown aerobically at 30°C in a shaking incubator (160 rpm) in TGY2X medium supplemented with the appropriate antibiotics. Typically for microscopy experiments, *D. radiodurans* cells were pre-grown the day before and then diluted for an overnight growth until reaching exponential (OD_650_ ∼ 0.3-0.5) the next morning. Optical density was measured on a Clariostar (BMG Labtech) plate reader. Ampicillin treatments were performed by growing cells at 30°C in medium containing 1 µg/mL ampicillin for 1 or 2h prior to PG labeling (see details below).

### Cell labeling for confocal and single-molecule localization microscopy (SMLM)

Membranes of exponentially growing *D. radiodurans* cells (1mL) were stained by addition of Nile Red (30 µM for confocal microscopy and 30-100 nM for SMLM) to the TGY2X growth medium for 10 min at room temperature. The cells were then harvested by centrifugation and resuspended in 200 μL TGY2X for confocal microscopy or instead washed 3 times in DPBS (3x 1 mL) and resuspended in 200 µL DPBS for SMLM. 3-5 µL of cell suspension was then deposited on a 1.5% (w/v) low melting agarose (LMA; Bio-Rad) pad prepared using a gene frame positioned on a glass slide. Two stripes of LMA were cut out on either side of the deposited sample for good aeration of the bacteria before a 1.5H coverslip was added to cover the frame. For PALM imaging, the DR-*FtsZ-mEos4B* strain was grown to exponential phase, washed twice with DPBS and deposited directly on a 1.5% LMA pad prepared in DPBS using a gene frame positioned on a glass slide. To visualize dividing bacteria in different orientations, an alternative set-up was also used (SI Fig. S5B) in which 10 μL of the cell suspension was placed on the bottom of a glass dish and cells were allowed to sediment for 2 min. Excess liquid was then gently removed using a pipette and after 2 min of air-drying, 10 µL 1.5% LMA equilibrated at 37°C was poured over the cells. The LMA was prepared in TGY2X medium for timelapse confocal imaging and in DPBS for PAINT imaging of Nile Red labeled bacteria. For PG labeling, 1.5 mL of exponential phase *D. radiodurans* cultures were centrifuged at 3,000 x *g* and resuspended in 200 µL TGY2X medium to which 50 µL of 10 mM azido-D-Alanine (aDA) was added^8,35^. Pulse labeling was typically performed for 10 min at 30°C, before washing the cells twice with cold DPBS to stop cell growth. Cells were then resuspended in 48 µL of 30 µM DBCO-AF488 or DBCO-AF647 (for dSTORM) diluted in DPBS and incubated on ice for 45 min to allow the DBCO-AF488/AF647 to enter the bacteria and react by click chemistry with the incorporated aDA. When ready to be imaged, labeled cells were washed twice with DPBS and resuspended in 100 µL of DPBS. For immunolabelling of HA-tagged FtsZ, the DR-*FtsZ-HA* strain (GY15705) expressing HA-tagged FtsZ was grown to exponential phase. 0.5 mL culture was flash-frozen using liquid nitrogen in DPBS-glycerol buffer (19% glycerol) supplemented with 2.58% formaldehyde. Cells were then fixed through slow thawing of these samples on ice overnight, washed twice in DPBS and resuspended in 100 µL of DPBS. Cells were permeabilized by treatment with 4 mg/mL lysozyme at 37°C for 30 min followed by the addition of 0.1% Triton X-100 for 5 min at 25°C. Cells were then washed twice with DPBS and incubated with a mouse anti-HA antibody (1:400 dilution in PBS-Tween 0.05% supplemented with 2% BSA) for 1 h at 37°C. After several washes with PBS-Tween 0.05%, cells were incubated with anti-mouse secondary antibody coupled to either AF488 (for confocal) or AF647 (for dSTORM) for 1 h at 37°C. After a final wash step, the bacteria were deposited on a 1.5% LMA pad prepared in DPBS as described above. For dSTORM experiments, the LMA was prepared in glucose buffer (62.5 mM Tris-HCl pH 8.0, 12.5% glucose, 12.5 mM NaCl) and contained 0.1M MEA and 1X GLOX (prepared from the 10X GLOX solution composed of 10 mM Tris-HCl pH 8.0, 50 mM NaCl, 56 mg/mL glucose oxidase and 13.6 mg/mL catalase). For confocal microscopy, the LMA was prepared as above in TGY2X.

### Confocal data acquisition and processing

Spinning-disk confocal microscopy was performed using an Olympus IX81 inverted microscope equipped with a Yokogawa CSU-X1 confocal head. The excitation laser beam (Ilas2 laser bench, GATACA systems) was focused to the back focal plane of a 100X 1.49-numerical-aperture (NA) oil immersion apochromatic objective. Series of Z-planes were acquired every 132 nm using a PRIOR N400 piezo stage to achieve cubic voxels. Fluorescence excitation was performed at 488 nm for AF488 and 561 nm for Nile Red. Fluorescence emission was collected with an Andor iXon Ultra EMCCD camera through a quad-band Semrock™ Di01-T405/488/568/647 dichroic mirror and single-band emission filters adapted to each fluorophore used: 520 nm for AF488 (FF02-520/28 Semrock™), and 600 nm for Nile Red (ET600/50m Chroma™). Data acquisition was performed using Metamorph 7.10 (Molecular devices). For Nile Red time-lapse imaging (SI Movies 1, 2 and 3), Z-stacks were acquired every 10 or 15 min. Acquired images were processed using Imaris (Oxford Instrument™) to correct for the possible translational (in x, y, and z directions) and rotational (z axis) drifts, followed by correction of the timepoints intensities using the embedded “Normalized timepoint” routine (Imaris XT package). Z-stacks of two-color images of Nile Red and PG-labeled *D. radiodurans* were deconvoluted in Imaris using 6 iterative cycles and calculated point spread functions (488 nm/520 nm and 561 nm/600 nm). When needed, Kymograph builder plugin in Fiji was applied to datasets. Coordinates of kymograph boundaries were then extracted in Fiji and used to calculate the septal closure rate in growing cells.

### SMLM (PAINT/dSTORM) data acquisition and processing

PAINT and dSTORM data were acquired on an Olympus IX83 inverted motorized microscope equipped with a SAFE 360 (Abbelight) SMLM set up, and using a UPLXAPO 100x oil immersion objective (NA 1.5, Olympus™). Data collection was performed using simultaneous dual camera acquisitions (50/50 splitter; Orca Fusion sCMOS – Hamamatsu™) with an astigmatism lens intercalated in the emission path of the direct camera to reconstruct 3D volumes in parallel with 2D single-molecule localization determination. Data was acquired at 27°C under continuous HiLo illumination with 400 W/cm² 561 nm light or 642 nm light and a typical frame time of 10 ms for PAINT and 50 ms for dSTORM. Typically, 40,000 – 60,000 frames were acquired per dataset under constant activation of the Zero Drift Control system (ZDC2 at 830 nm) to limit Z-drift during acquisitions, and at constant temperature (Digital Pixel™ blind cage incubator and water jacket around the objective set at 27°C). Typical 3D reconstructions are limited by the astigmatic point spread function to a few hundreds of nm. When specified in the corresponding figure legends, we collected stacked localizations at 400 nm distance and reconstructed the cumulated localizations to obtain larger reconstructed volumes. SMLM data was processed using NeoAnalysis software (Abbelight™) using default values for maximum likelihood estimations (MLE) of the gaussian localizations fittings. Further filtering of the dataset was performed using Thunderstorm plugin^66^ in Fiji^67^ to correct for possible drift in x and y directions (using cross-correlation), and restrict the localizations to limit the spreading of values for sigma, intensity, and uncertainty values.

### Sample preparation, vitrification and cryo-FIB milling for cellular cryo-ET

A 20 mL pre-culture of *D. radiodurans* cells was grown overnight to stationary phase (OD_650_ ∼ 1.8) in TGY2X medium in a shaking incubator at 30°C and 170 rpm and then diluted to OD_650_ ∼ 0.1 in 20 ml fresh medium the next day and grown further until reaching exponential phase (OD_650_ ∼ 0.4). The culture was then centrifuged for 5 min at 4,000 rpm, the pellet collected in a 1 mL Eppendorf tube and washed three times in 1 mL PBS. Shortly before plunge-freezing, the pellet was resuspended in PBS to reach a final volume of ∼ 60 μL. Plunge-freezing was performed in liquid ethane/propane mixture using Vitrobot Mark IV (Thermo Fisher Scientific) at 23°C, 90% humidity, with blot force −5 to 10, blot time 8 to 10 sec and wait time 30 sec. 4 μL of cell suspension were deposited per Quantifoil Cu 1.2/1.3, 200 mesh grid (Micro Tools GmbH, Großlöbichau, Germany), glow-discharged immediately prior to use. Frozen grids were clipped into standard AutoGrid specimen cartridges (Thermo Fisher Scientific) marked to keep track of the milling direction for the subsequent orientation in the Titan Krios (Thermo Fisher Scientific) microscope, and stored in liquid nitrogen until usage. For cryo-FIB milling, the clipped grids were mounted into a 45° pre-tilt shuttle and transferred into a Scios cryo-FIB/scanning electron microscope dual-beam microscope (Thermo Fisher Scientific). To reduce curtaining and enhance sample conductivity, the grids were sputter-coated with organometallic platinum using the gas injection system. The Gallium milling was performed at a 17° to 23° stage tilt angle, dependent on the milling location on the grid. Lamellae were prepared in a stepwise manner, progressively reducing the FIB current from 0.5 nA to remove the bulk material to 30 pA for the final polishing step, ending up with lamellae of 200-300 nm thickness. Progress of the milling process was monitored using the SEM operated at 10 kV and 50 pA. Grids were stored in liquid nitrogen until transfer into the Titan Krios microscope.

### Cryo-ET data acquisition

Data was collected using a Titan Krios operated at 3001keV and equipped with a BioQuantum post-column energy filter (Gatan, Pleasanton, CA) and a K2 Summit direct detector (Gatan, Pleasanton, CA). The energy filter was operated with a 201eV slit width. Image acquisition was performed using SerialEM software. To identify regions of interest (ROI) and account for the lamella pre-tilt, lamella montages were acquired at an intermediate magnification of 17 Å/pixel and at +13° tilt. Tilt series were then recorded with the detector operated in super-resolution mode, at a nominal magnification of 33,000x, corresponding to 2.1731Å/pixel at the specimen level. Data were collected at ∼ 3 µm defocus following a grouped dose-symmetric tilt scheme in 3° increments, with a tilt range depending on the ROI position and thickness, typically ± 50° to ± 60°. The dose was kept constant for all tilts in each given tilt series and adjusted such as to reach the total per tilt series dose of ∼ 140 e/Å^2^. In total, 45 tilt series were collected for the presented analysis.

### Cryo-ET image processing, visualization, segmentation and analysis

Data preprocessing was carried out in Warp^68^. Gain reference and motion correction, and CTF estimation were performed following the standard Warp^68^ procedure for tomography data with the following non-default parameters: binning 1x, CTF grid dimensions 5x5x1, and motion grid dimensions 5x5x15. The dataset was then manually curated to remove low quality tilt images and extracted into stacks for tilt series alignment. Alignment was performed in batch with AreTomo^69^, using our in-house script Waretomo (<u>source code</u>) to integrate it seamlessly with Warp^68^. Alignment parameters were reimported in Warp^68^ and used to reconstruct full tomograms at a resolution of 17.41 Å/pixel. Tomograms were denoised using Topaz^70^ to help with picking and inspection. All images of tomogram slices presented in this manuscript come from non-denoised tomograms, averaging over 5 slices. Cell walls were annotated using the surface annotation tool in blik^32^ (<u>source</u> <u>code</u>). These annotations were then used to generate the 3D surface visualizations of the septa and cell wall profiles using the surface tool in blik^32^, resampling the tomogram perpendicularly to the surfaces to obtain ‘straightened’ wall volumes. 2D projections were obtained by averaging over the Z dimension of the resampled volumes. 1D density profiles were calculated in Fiji^67^ on 76-pixel sections of the 2D projections and used to measure cell wall dimensions using the point to point measuring tool in Fiji^67^ as detailed in SI Fig. S8.

## ACKNOWLEDGEMENTS

IBS acknowledges integration into the Interdisciplinary Research Institute of Grenoble (IRIG, CEA). The optical imaging was carried out on the M4D imaging platform of the Grenoble Instruct-ERIC center (ISBG; UAR 3518 CNRS-CEA-UGA-EMBL) within the Grenoble Partnership for Structural Biology (PSB), supported by FRISBI (ANR-10-INBS-0005-02) and GRAL, financed within the University Grenoble Alpes graduate school (Ecoles Universitaires de Recherche) CBH-EUR-GS (ANR-17-EURE-0003). We thank staff on the M4D platform at IBS for their help for data acquisition and processing, and the Abbelight team for help with data acquisition. Electron microscopy sample preparation, cryo-FIB milling and cryo-ET data collection were performed at the Umeå Centre for Electron Microscopy (UCEM), the Umeå Cryo-EM node of SciLifeLab national research infrastructure. IG thanks the UCEM staff for training and support.

