## Supplemental videos for "Combining live cell fluorescence imaging with *in situ* cryo electron tomography sheds light on the septation process in *Deinococcus radiodurans*"

### Slide 1
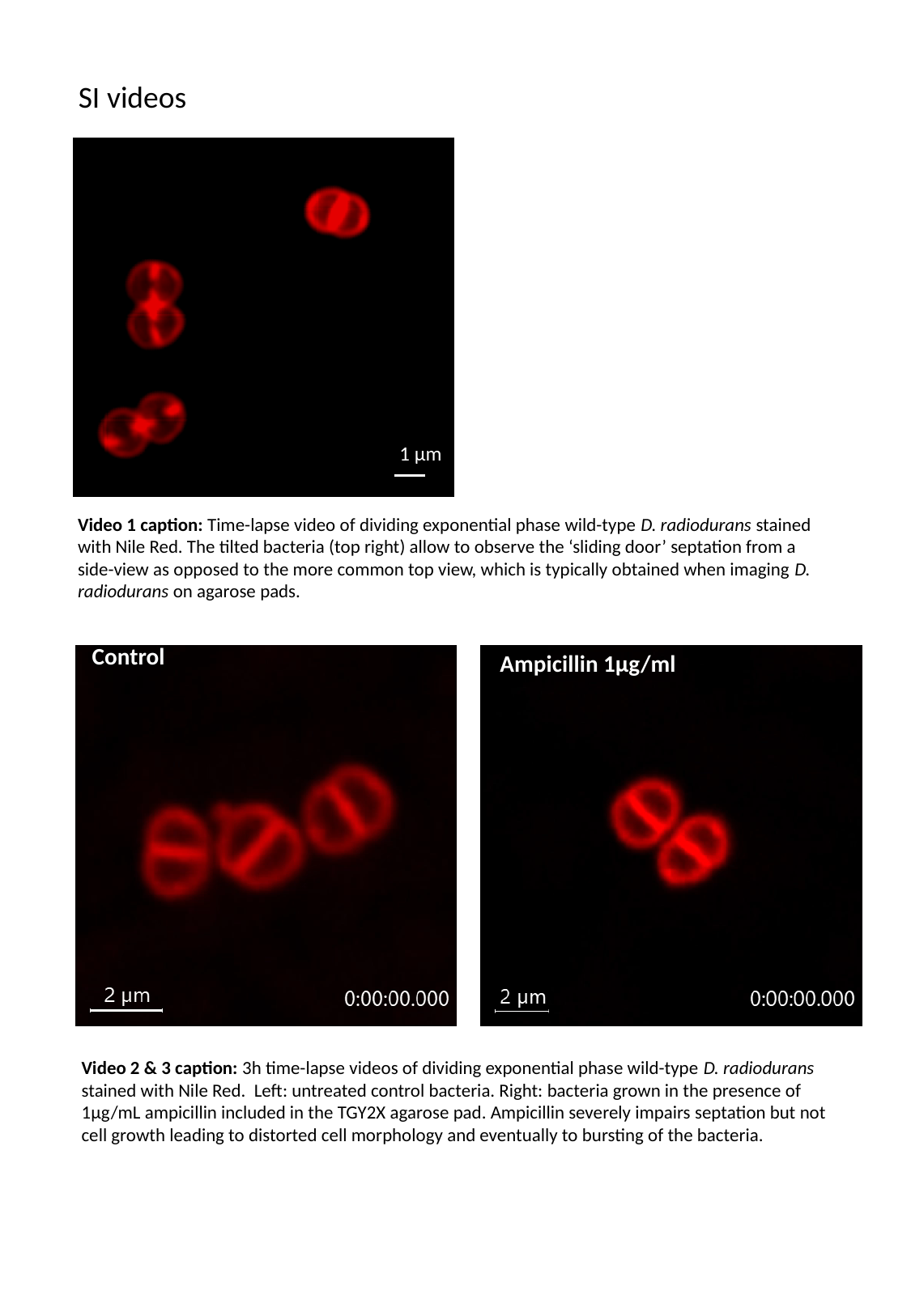

SI videos
1 µm
Video 1 caption: Time-lapse video of dividing exponential phase wild-type D. radiodurans stained with Nile Red. The tilted bacteria (top right) allow to observe the ‘sliding door’ septation from a side-view as opposed to the more common top view, which is typically obtained when imaging D. radiodurans on agarose pads.
Control
Ampicillin 1µg/ml
Video 2 & 3 caption: 3h time-lapse videos of dividing exponential phase wild-type D. radiodurans stained with Nile Red. Left: untreated control bacteria. Right: bacteria grown in the presence of 1µg/mL ampicillin included in the TGY2X agarose pad. Ampicillin severely impairs septation but not cell growth leading to distorted cell morphology and eventually to bursting of the bacteria.
