## Supplemental figures S1-S10 for "Combining live cell fluorescence imaging with *in situ* cryo electron tomography sheds light on the septation process in *Deinococcus radiodurans*"

### Supplementary figures

#### SI Fig. S1

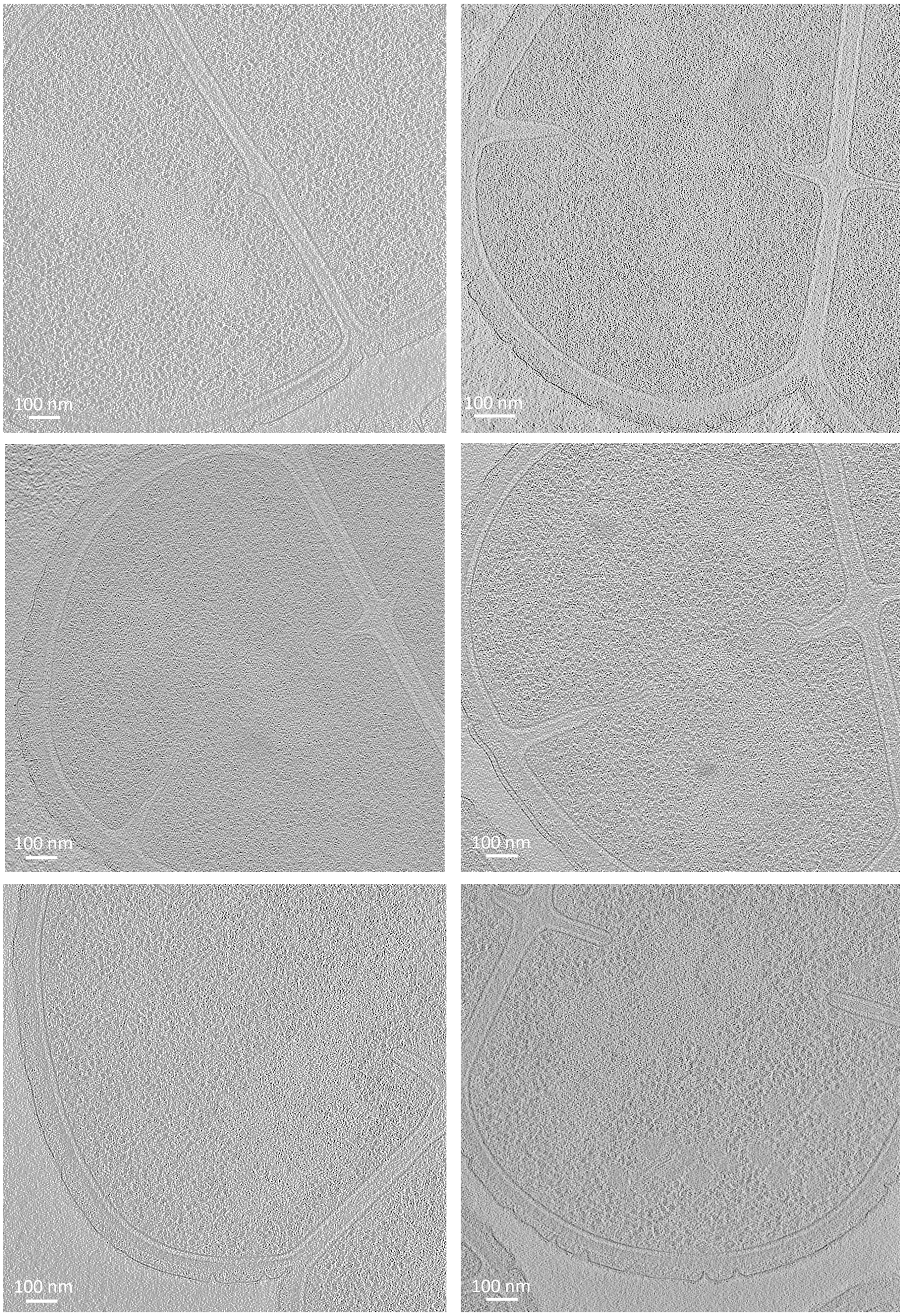


**Figure S1**: Examples of 2D slices through selected tomograms collected on early-phase dividing *D. radiodurans* cells.

### SI Fig. S2

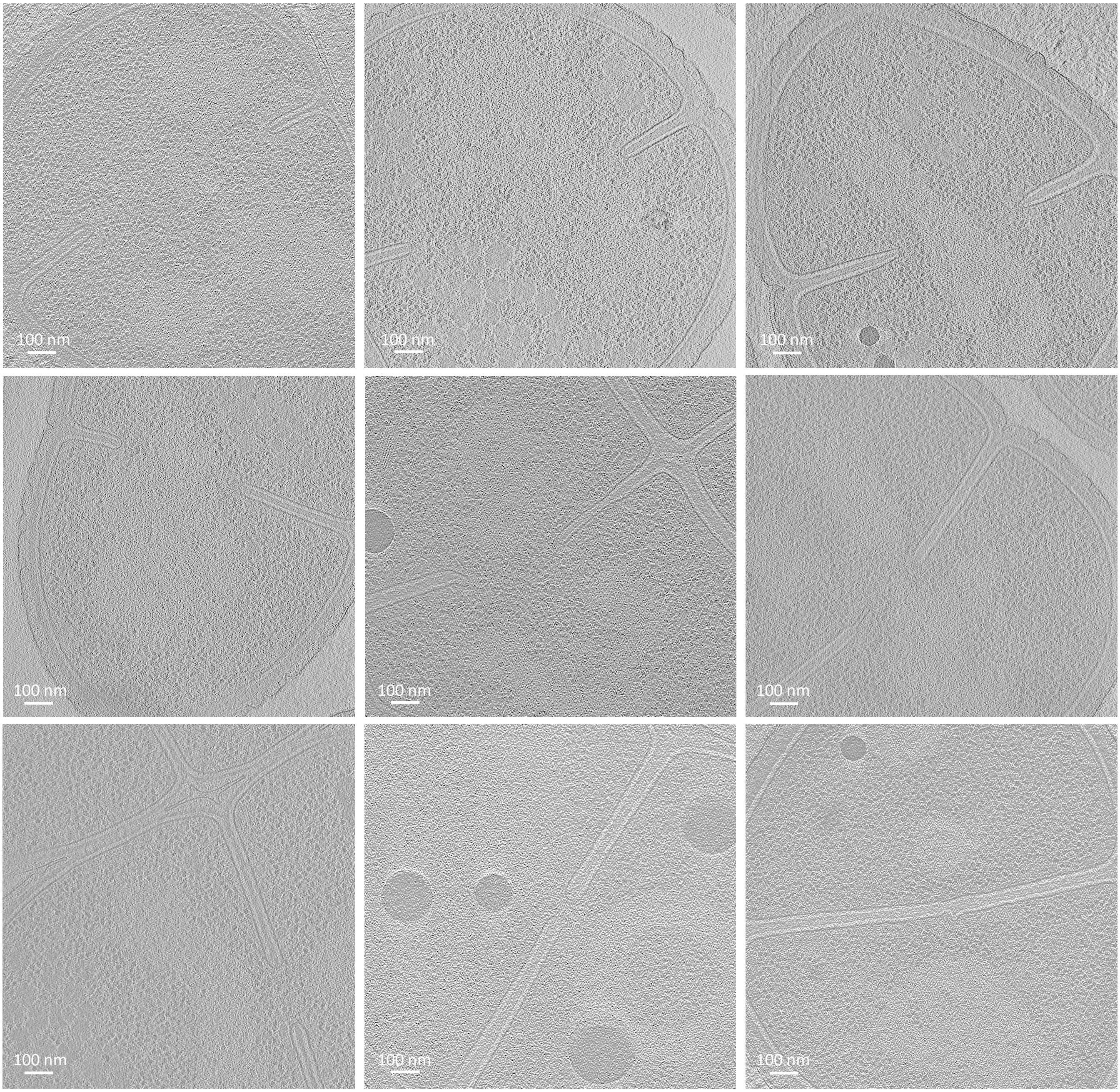


**Figure S2**: Examples of 2D slices through selected tomograms collected on mid- to late-phase dividing *D. radiodurans* cells.

### SI Fig. S3

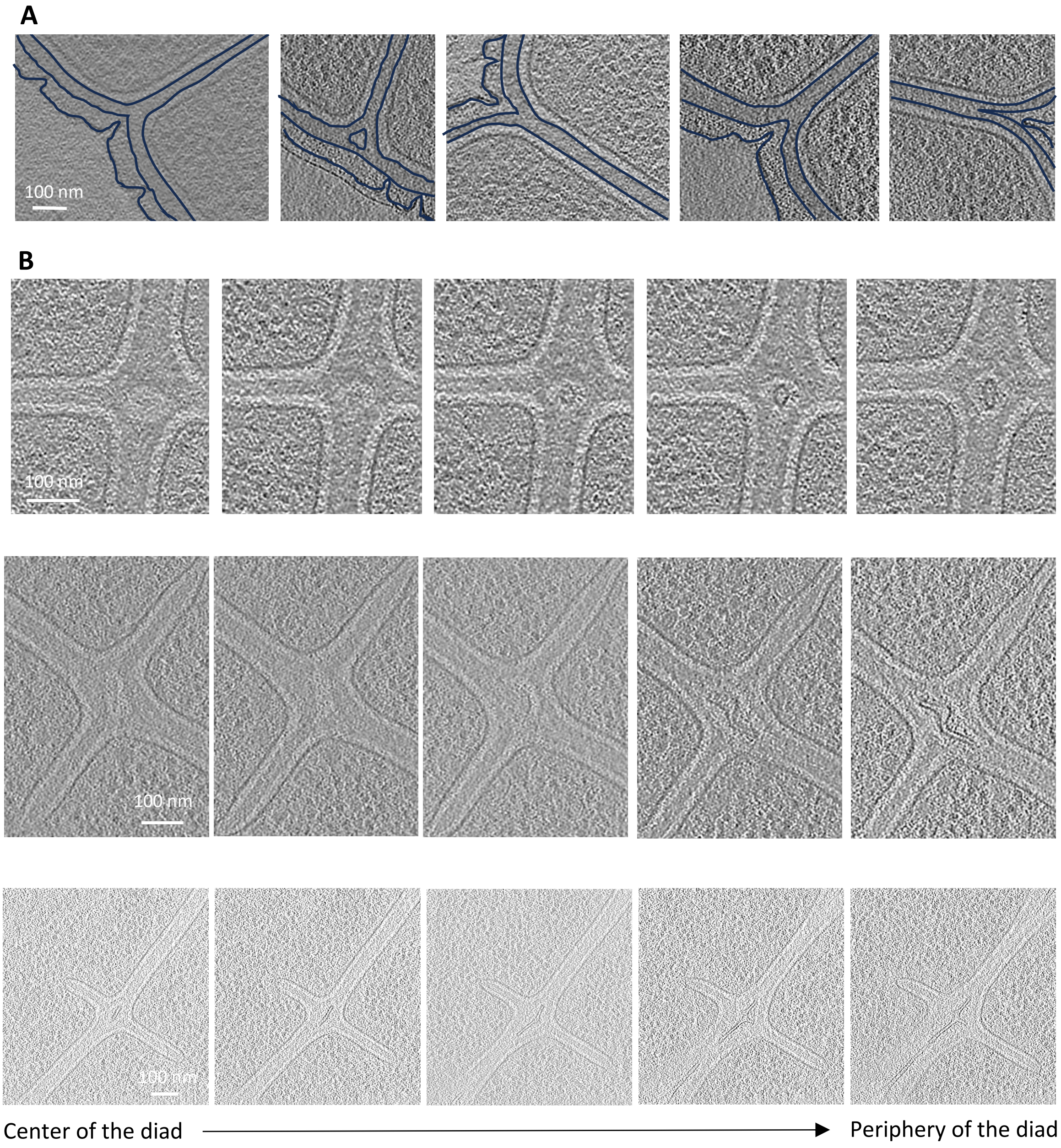


**Figure S3**: (A) Examples of junctions between the peripheral cell envelope and the growing septa observed in various tomograms of *D. radiodurans.* The borders of the different layers composing the cell wall are highlighted in dark blue. (B) Examples of regions in which splitting of the daughter cells were observed. From left to right: Z-slices through the tomograms going from the center of the diads (left) to the outer periphery of the bacteria (right).

### SI Fig. S4

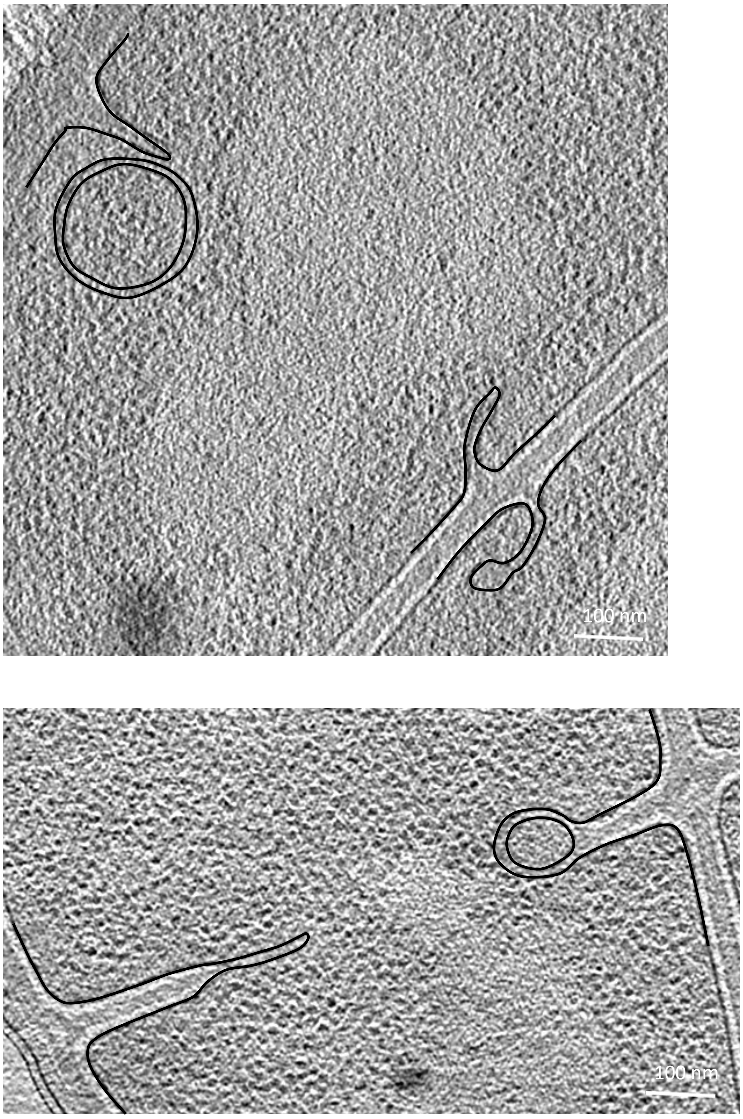


**Figure S4**: Examples of membrane protrusions observed in various tomograms of *D. radiodurans* at the leading edge of growing septa. The IM is highlighted in black.

### SI Fig. S5

**A**

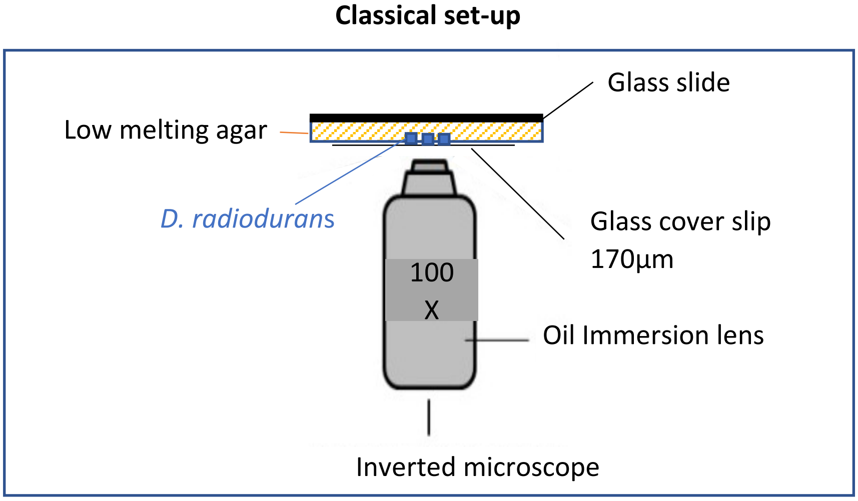


**B**

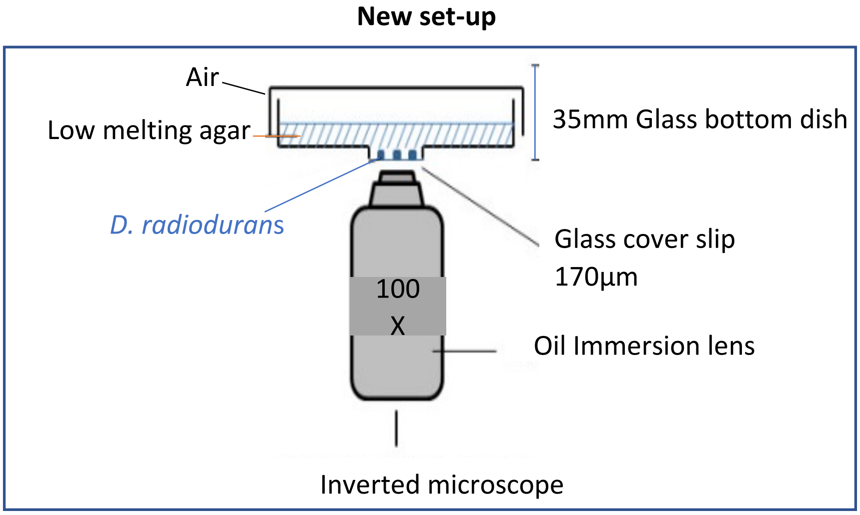


**Figure S5**: Microscopy set-ups for timelapse 3D confocal video-microscopy. (A) Classical set-up in which cells are seeded on a low melting agarose pad and then overlaid with a glass coverslip. In this set-up, cells are mostly positioned in the same orientation orthogonal to the two successive division planes. (B) New set-up in which cells are deposited on the glass of a glass-bottomed dish and then coated with low melting agarose. The pouring of the melted agarose over the cells results in a wider distribution of cell orientations allowing to capture side or face views of the septation in tilted cells.

### SI Fig. S6

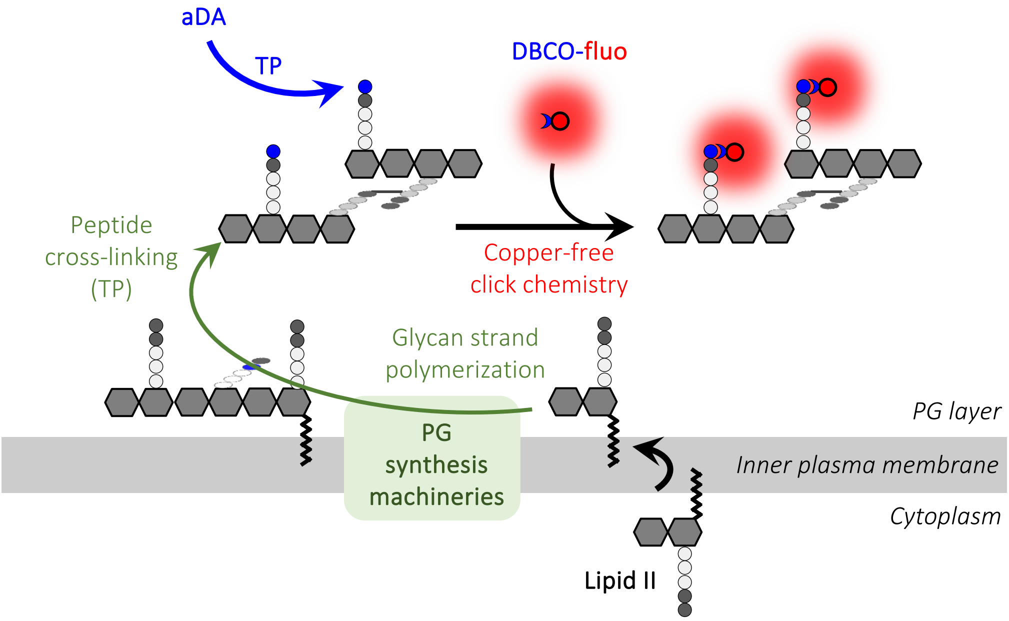


**Figure S6**: Schematic diagram of the incorporation of the azido-D-Alanine (aDA) probe by transpeptidases (TP) into the peptide chains of the peptidoglycan (PG). The incorporated probe is revealed by its conjugation, via copper-free click chemistry, to a fluorescent dye (Alexa Fluor 488 or Alexa Fluor 647) carrying a dibenzocyclooctyne (DBCO) group.

SI Fig. S7

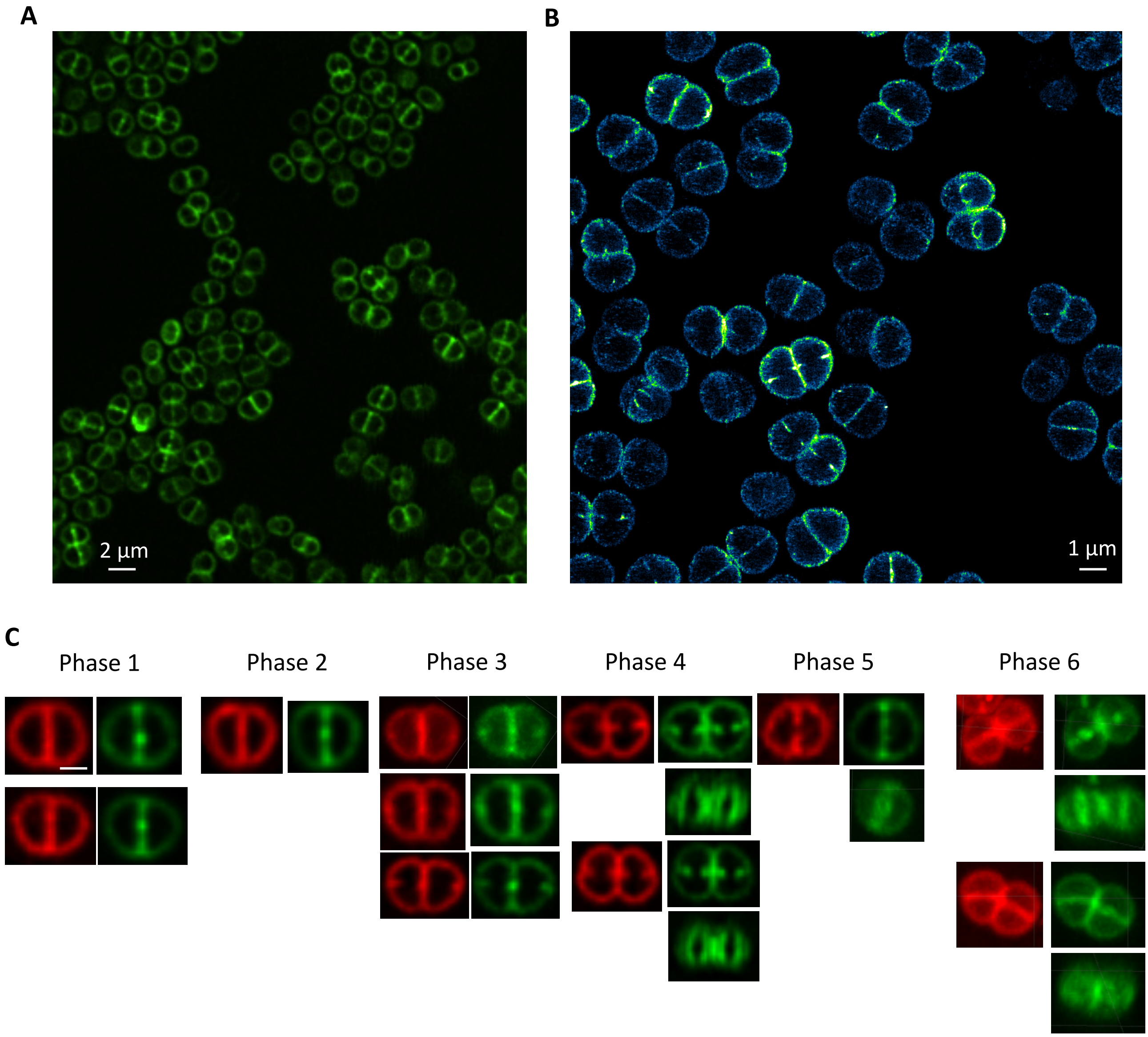


**Figure S7**: Examples of images of PG-labeled *D. radiodurans* cells viewed by confocal (A) and dSTORM (B) microscopy. (C) Dual labelling of PG (green) and membrane (Nile Red) layers as a function of the phases of the cell cycle. Scale bar = 1 µm.

SI Fig. S8

**A**

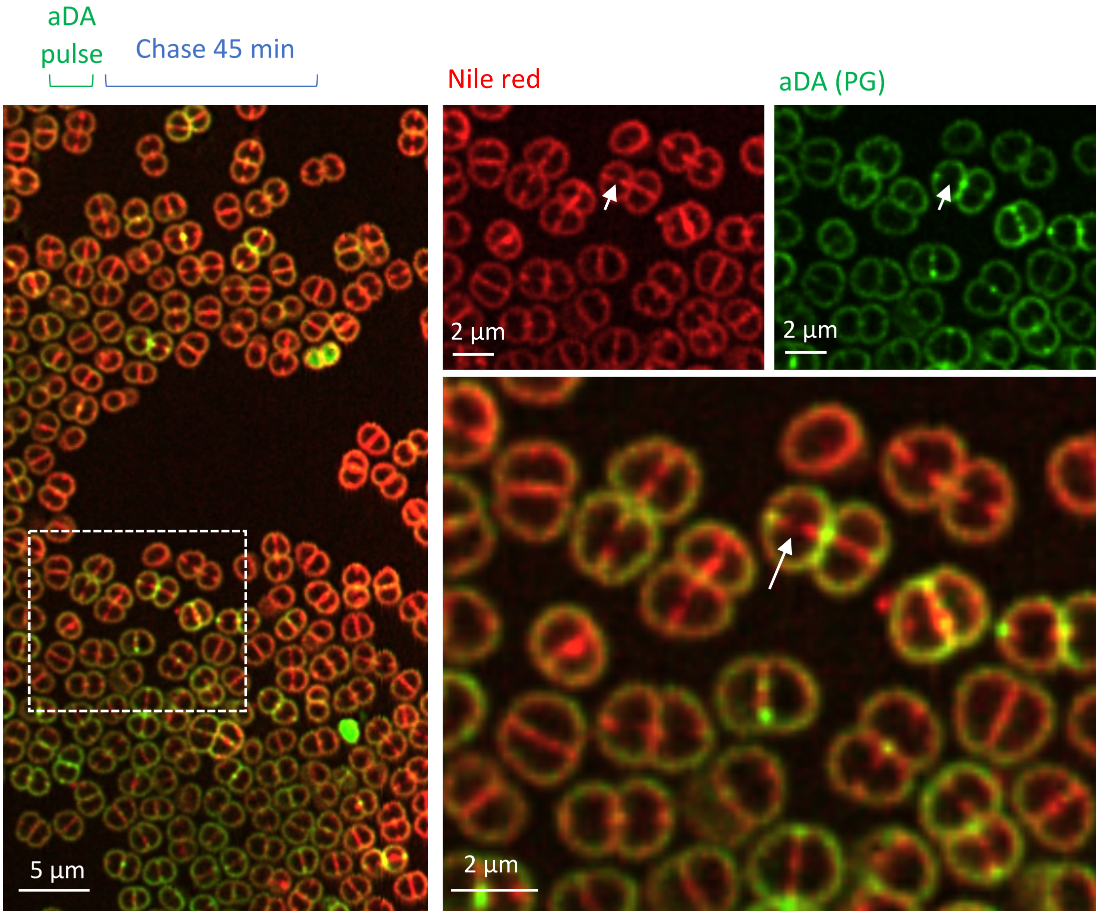


**B**

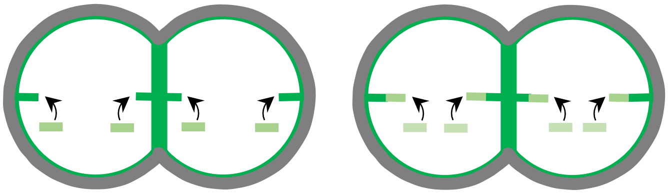

**Figure S8**: (A) Examples of deconvoluted two-color confocal microscopy images (single Z-slice) of PG (green) and Nile Red (red) labeled *D. radiodurans* after a 45 min chase experiment (left). The right panels correspond to a close-up view of the region boxed in white in the left panel. Top right: split green and red channels. Bottom rigt: overlaid green and red channels. White arrows indicate Nile Red labelled growing septa that have largely lost their aDA PG labelling (green) after the chase period. (B) Diagram illustrating the stepwise synthesis of PG at the tip of the growing septa to progressively extend the septa until they are sufficiently close to fuse.

SI Fig. S9

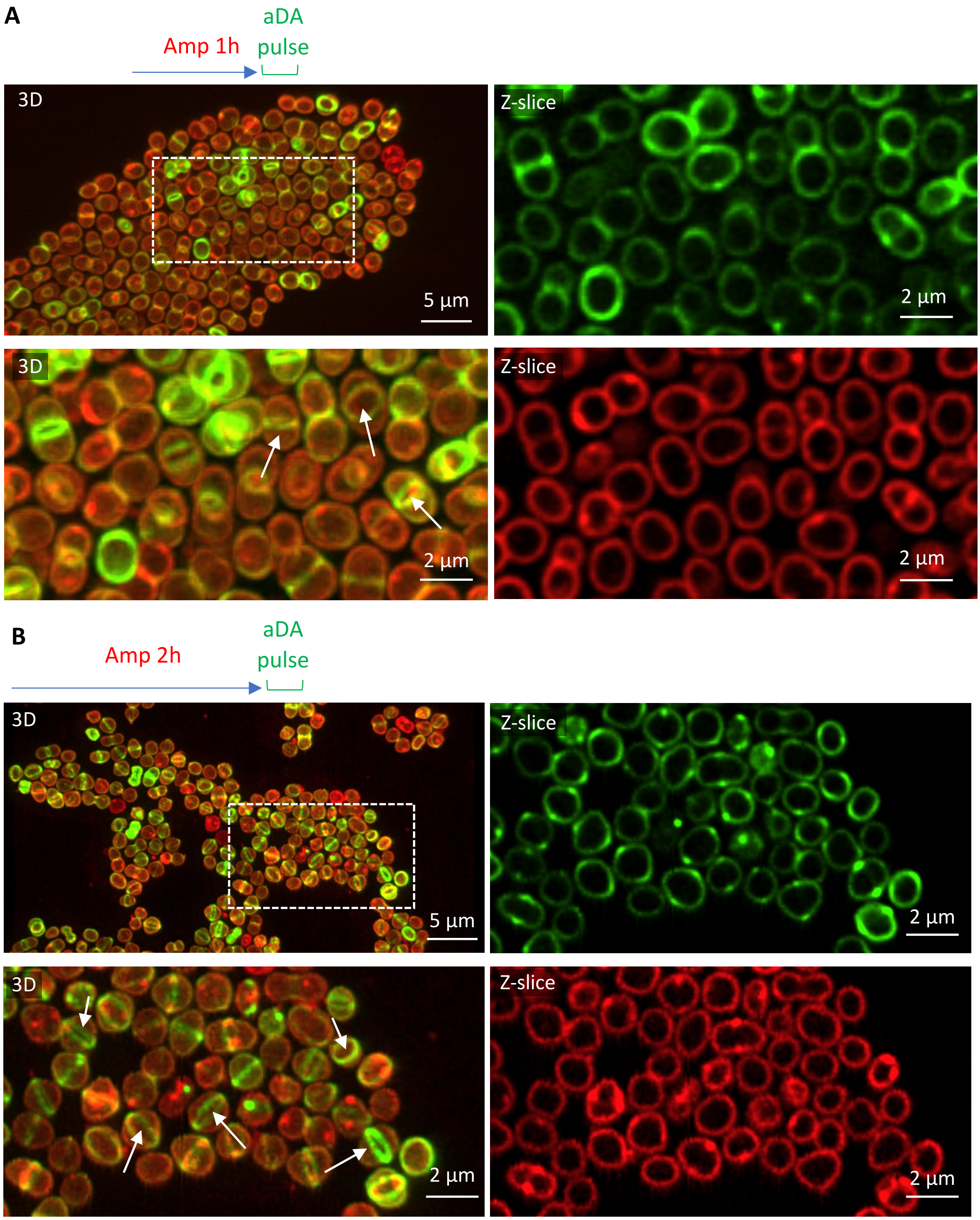


**Figure S9**: Examples of deconvoluted two-color confocal microscopy images of aDA-pulse labeled PG (green) and Nile Red (red) stained *D. radiodurans* after growth in the presence of 1 µg/mL Ampicillin for 1 h (A) or 2 h (B). (A-B) Top left: full 3D views. Bottom left: close-up 3D views of the regions boxed in white in the top left panels. Right: close-up 2D views (single Z-slices) of the boxed region with split green (top) and red (bottom) channels. White arrows indicate ring-shaped aDA labeling (green) located at the division site.

### SI Fig. S10

#### Peripheral cell envelope

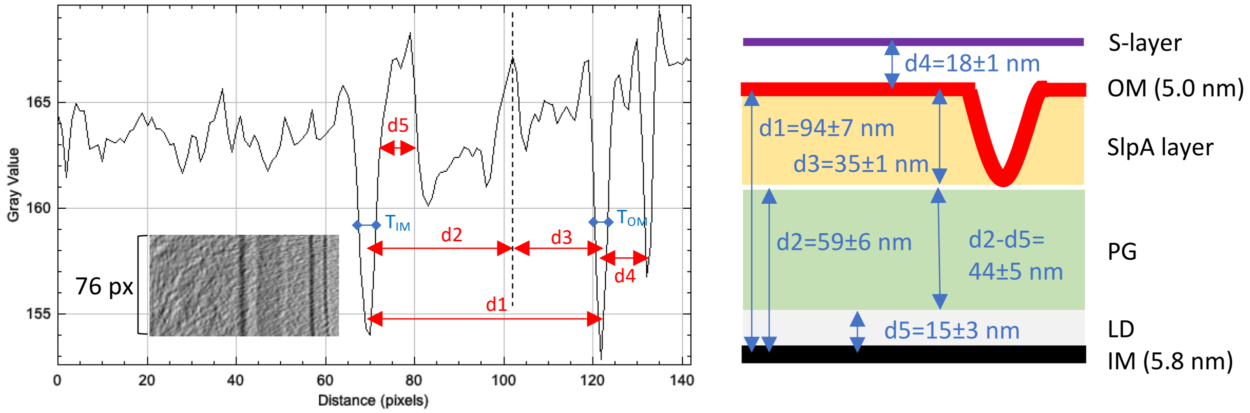


#### 'Old' septum

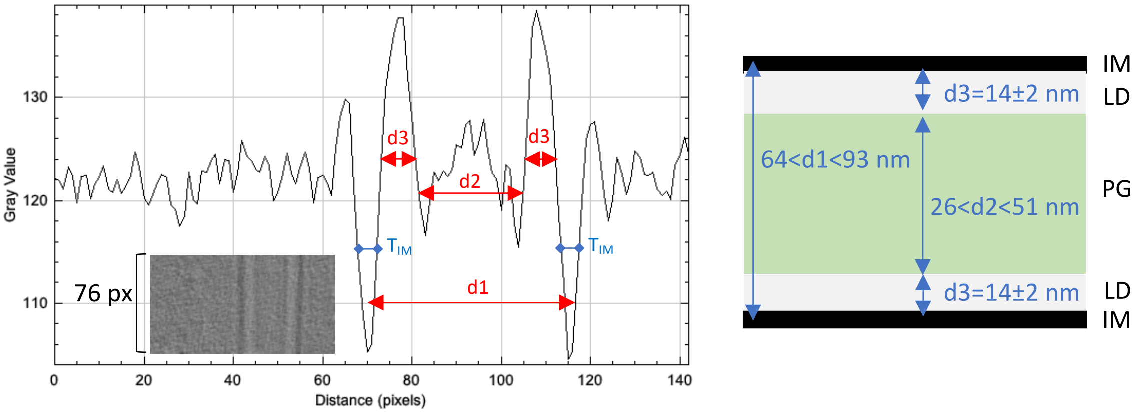


#### Growing septum

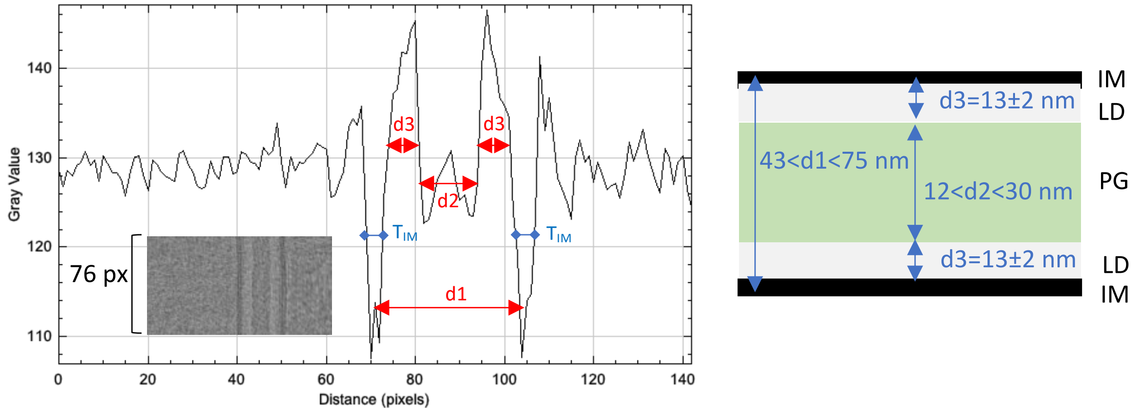


**Figure S10**: Typical 1D density profiles obtained for 76-pixel sections of 2D projections of straightened cell walls (illustrated as insets) from the peripheral cell envelope (top), 'old' septum (middle) and the growing septum (bottom). Various measurements were made to determine the average thicknesses of the different layers composing these distinct cell wall regions. Distances between the middle of the IM, OM, central white line and the S-layer were measured as shown by the red arrows. Additionally, the thickness of the inner ($T_{IM}$) and outer ($T_{OM}$) were measured at mid-peak height as shown with the blue lines. The thickness of the periplasmic space (d5 in top panel and d3/d3' in lower panels) was measured at the base of this positive peak.
